# Seasonal variation in insect assemblages at flowers of Balanites aegyptiaca, an ecologically and socially important tree species in the Ferlo region of Senegal’s Great Green Wall corridor

**DOI:** 10.1101/2025.03.26.645493

**Authors:** Natalia Medina-Serrano, Anne-Geneviève Bagnères, Mouhamadou Moustapha Ndiaye, Valentin Vrecko, Doyle McKey, Martine Hossaert-McKey

**Affiliations:** Centre d’Ecologie Fonctionnelle et Evolutive (CEFE), CNRS, Univ Montpellier, EPHE, IRD, Montpellier 34293, France; Institut Fondamental d’Afrique Noire Cheikh Anta Diop (IFAN Ch. A. Diop), Université Cheikh Anta Diop (UCAD) de Dakar, Senegal

**Keywords:** *Balanites aegyptiaca*, biotic interactions, flower-visiting insects, Great Green Wall, insect-tree interactions, restoration, Sahelian ecosystems, pollination

## Abstract

Interactions between flowers and flower-visiting insects play central roles in ecosystem functioning. In addition to ensuring pollination, flower-visiting insects are also crucial for numerous other biotic interactions, using floral resources as fuel in their search for prey, hosts, breeding sites, and other resources. Studying insect-flower interactions may thus be strategic to conserve and restore biotic interactions in ecosystems heavily degraded by intensive land use and climate change, such as the arid savannas of the northern Sahel. We aimed to document for the first time the diversity of flower-visiting insects in this region and to examine whether restoration efforts of the Great Green Wall initiative have affected insect abundance and diversity. Using two capture methods, hand netting and pan traps, we inventoried insects visiting flowers of *Balanites aegyptiaca*. This is the most abundant tree species in the region and is also of economic importance. We sampled three sites in a single locality: a “Restored” site from which livestock are excluded, an ”Unrestored” site in intensively grazed rangeland, and a topographical ‘“Depression” site in a grazed area but with high tree density. Each site was sampled at three different periods to examine variation in this extremely harsh and seasonal environment. The open-access flowers of *Balanites aegyptiaca* are produced in multiple flowering events each year. We found a surprisingly high diversity of insects visiting its flowers, comprising 371 morphospecies from 10 insect orders, with a predominance of Hymenoptera and Diptera. Insect abundance, diversity and species composition differed markedly between seasons. Bees appear to be important pollinators, particularly solitary bees of the family Halictidae, which were abundant in all seasons. Diptera, in particular several families of small flies, were also frequent visitors and were especially abundant and diverse in the wet season. Ants were especially abundant at flowers in the dry season, when few resources other than flowers and flower-visiting insects were likely available to them. Insect abundance and observed diversity differed only little between sites, but estimated total diversity (Chao1 richness) was highest in the Depression site. Insects visiting flowers of *B. aegyptiaca* included herbivores, decomposers, and parasitoids and predators of diverse arthropods, underlining the role of these floral resources in numerous facets of ecosystem functioning. The floral resources of *B. aegyptiaca* and other trees, which can flower throughout the year, are likely critical to assure the persistence of numerous insect species. Integrating biotic interactions into ecosystem management is crucial for conservation and restoration in Sahelian ecosystems.

## Introduction

The Sahel is among the world’s regions most vulnerable to climate change. In the semi-arid savannas of the northern Sahel, the combination of climate change and land-use change, particularly intensive grazing and browsing, has led to widespread environmental degradation. Initiatives such as the United Nations Convention to Combat Desertification (UNCCD) and the pan-African “Great Green Wall” (GGW) attempt to halt and reverse this degradation, with varying degrees of success (Delay et al., 2022; Turner et al., 2023). With recent calls to “accelerate” these initiatives (UNCCD, 2021; Diallo et al., 2023), it is essential to establish a more solid foundation for integrative approaches to ecosystem restoration. Work on restoration in the region has been plant-centred, with little attention to biotic interactions, which are crucial for ecosystem functioning and resilience (Carlucci et al., 2020; Dicks et al., 2021). A comprehensive approach to restoring pastoral ecosystems in the northern Sahel is currently lacking. Research has primarily focused on sustaining the productivity of silvopastoral systems, with relatively little attention to the impact of management options on biodiversity (e.g., Touré et al., 2012; Rasmussen et al., 2018). Revegetation actions of the GGW, principally planting trees and encouraging their natural regeneration (Dia & Duponnois, 2012; Mugelé, 2020), can result in improved pollination services by increasing the amount and diversity of floral resources in space and time, but maximising their impact depends on selecting appropriate species, based on knowledge about how different tree species interact with flower-visiting insects (Kireta et al., 2024). Like other insects, flower-visiting species also require food resources for immature stages, which often feed at trophic levels different from adults, and suitable habitat for each stage in their complex life cycles (Buisson et al., 2024). Understanding biotic interactions is crucial for developing actions to maintain and restore biodiversity in the Sahel, especially given concerns that decreasing habitat connectivity could lead to a loss of rare species and of particular ecosystems (Menz et al., 2011). Our research focuses on insect-flower interactions in the context of theGGW initiative in the Ferlo region of northern Senegal. This focus provides an entry point to explore the diversity of biotic interactions and their contributions to ecosystem functions (Kevan, 1999). Insects play a crucial role in the pollination of angiosperms, making them key players in ecosystems, and maintaining their interactions with flowering plants is essential for successful ecological restoration (Dirzo et al., 2014; Genes & Dirzo, 2022). Floral resources are also crucial for many insects that may not be effective pollinators but that contribute to other ecosystem functions such as population regulation, decomposition and nutrient cycling (Medina-Serrano et al., 2025).

The northern Sahel is characterised by a long and harsh dry season. How does seasonality affect insect-flower interactions? Whereas some insects remain active year-round, adult stages of most insects in seasonally dry tropical regions show marked seasonal variation. Furthermore, the activity of some insects peaks in the wet season, while others peak in the dry season (Kishimoto-Yamada & Itioka, 2015). Many studies of insect seasonality have compared rain forests and tropical dry forests (e.g., Janzen & Schoener, 1968; Kishimoto-Yamada & Itioka, 2015) or examined differences in seasonal patterns between wetter and drier sites within tropical dry forest (Janzen & Schoener, 1968). Fewer studies have been done in savanna (Silva et al., 2011, Gaona et al., 2025), and often concern only certain insect groups such as termites (Davies et al., 2015) or ants (Leal & Oliveira, 2000). To the best of our knowledge there have been no community-level studies of insect seasonality in semi-arid Sahelian savannas.

The entomological literature for the Ferlo region, and for the northern Sahel in general, is scattered across various specialist publications. Mirroring the situation for much of Africa, a comprehensive synthesis on insect diversity is lacking, particularly concerning insects involved in pollination (Rodger et al., 2004; Archer et al., 2014). A primary objective of our research is thus to characterise the diversity of flower-visiting insects in the Ferlo region over the annual cycle and assess their current status, an essential step in studying the impact of GGW actions on ecological restoration (Medina-Serrano et al., 2025). The present paper focuses on insects that visit the flowers of *Balanites aegyptiaca* (L.) Delile (the desert date; Zygophyllaceae), the most common woody species in the studied region. With more than three flowering periods each year, the timing and duration of which vary across years, it is a major resource for flower-visiting insects. In addition to its ecological importance (Hall, 1992), this species supplies local people with fodder for livestock, edible fruits, and materials for construction (Sagna et al., 2014; Ousmane et al., 2023; Newby, 2024). To understand how the region’s extreme seasonality affects insect assemblages, we sampled the diversity of insects on flowers of this species in three different seasons: end of the dry season (June 2022), rainy season (August 2022) and middle of the dry season (February 2023). We also compared assemblages of flower visitors in three sites differing in their management history (either benefiting or not from GGW restoration actions aimed at restoring ecosystem functioning and biodiversity) and in tree density. By focusing on *Balanites aegyptiaca*, an ecologically and socially important tree species in the Sahel, we aim to fill crucial knowledge gaps related to insect-flower interactions in the region and their role in ecosystem functioning and provision of ecosystem services (Ndong et al., 2015; Dendoncker et al., 2020).

Specifically, we ask: How does seasonality influence the abundance, diversity, and composition of assemblages of flower-visiting insects on *Balanites aegyptiaca*Do restoration actions (in this case, exclusion of livestock) influence these parameters? Are there differences in insect assemblages between well-drained sites that dominate the landscape and scattered topographical depressions, in which tree density and diversity are much higher? To address these questions, we compared the diversity and composition (both taxonomic and functional) of flower-visiting insects across seasons and sites. We present (1) an overall analysis of the taxonomic and functional diversity and abundance of flower-visiting insects, (2) an assessment of the effects of seasonality and habitat management on these communities, and (3) a detailed evaluation of the seasonal patterns in assemblages of flower-visiting insects.

## Material and method

### (1) Study area

The study was conducted in the Ferlo region, northern Senegal, characterised by a semi-arid climate with average annual rainfall ranging from 300 to 500 mm, average temperature of approximately 28°C, and strong seasonality, with a 2-3-months rainy season and 9-10 months without any rain (Cissé et al., 2016). To monitor variation in weather conditions throughout the period of the study, we installed a sensor monitoring temperature and humidity every 30 minutes, and a rain gauge, in the village of Koyli Alpha, about 1 km from the closest sampling site and 6 km from the furthest. Figure S1 shows the distribution of precipitation during our study period. Figure S2 shows that mean temperature does not vary much throughout the year, but that large diurnal differences in temperature and humidity were registered during the dry season. This variation is produced by the Harmattan, a cool dry wind from the northeast or east in the western Sahara that is strongest from late November to mid-March. Its arrival may cause air temperatures to fall to 9 °C. One of the largest diurnal variations measured was in December 2022, from 89.5% relative humidity and 17.7 °C at 7:00 am, to 19.5% relative humidity and 33 °C the afternoon of the same day.

The Ferlo is dominated by herbaceous plants, particularly grasses (Poaceae) and leguminous forbs (Fabaceae). Vegetation in the Ferlo is classified as tree/shrub savanna, in which Balanites aegyptiaca, Vachellia tortilis subsp. raddiana (Savi) Kyal. & Boatwr., and Senegalia senegal (L.) Britton (the two last species are both Fabaceae: Mimosoideae) are the most abundant woody species. Other common woody species in the area include Boscia senegalensis (Pers.) Lam. ex Poir. (Capparaceae) and Calotropis procera (Aiton) W.T. Aiton (Apocynaceae). Vegetation in the northern Ferlo is subjected to intensive grazing and browsing by cattle, sheep, and goats. Degradation of soils and desertification (i.e., irreversible degradation of ecosystems), owing to the combined effects of climate change and human land use (mostly pastoralism in this region), are major environmental concerns (Miehe et al., 2010; Vincke et al., 2010).

The study area was located near the village of Koyli Alpha (Figure 1), situated in the southern part of the valley of the “Lac de Guiers”. Within this locality, we studied three sites. The site characterised as “Depression” is not within the restored area and is thus subjected to grazing and browsing by livestock, but is located in a topographical depression. In such lower-lying areas, soils retain moisture into the dry season and tree density and diversity are higher than in the two other sites (Dendoncker & Vincke, 2020; Dendoncker et al., 2023). The two other sites are comparable in terms of soil and drainage but differ in intensity of grazing and browsing. The “Restored” site is within the Koyli Alpha Community Natural Reserve, where livestock have been effectively excluded from an area of around 650 ha since 2017. Within the reserve, herders are allowed to cut and export herbaceous biomass at the end of the dry season. As most herbaceous plants in the region are annuals, this harvest occurs after production and dispersal of seeds and does not cause mortality of adult plants. The third site (“Unrestored”) is located in intensively grazed and browsed rangeland nearby (Figure 1).

**Figure 1.**
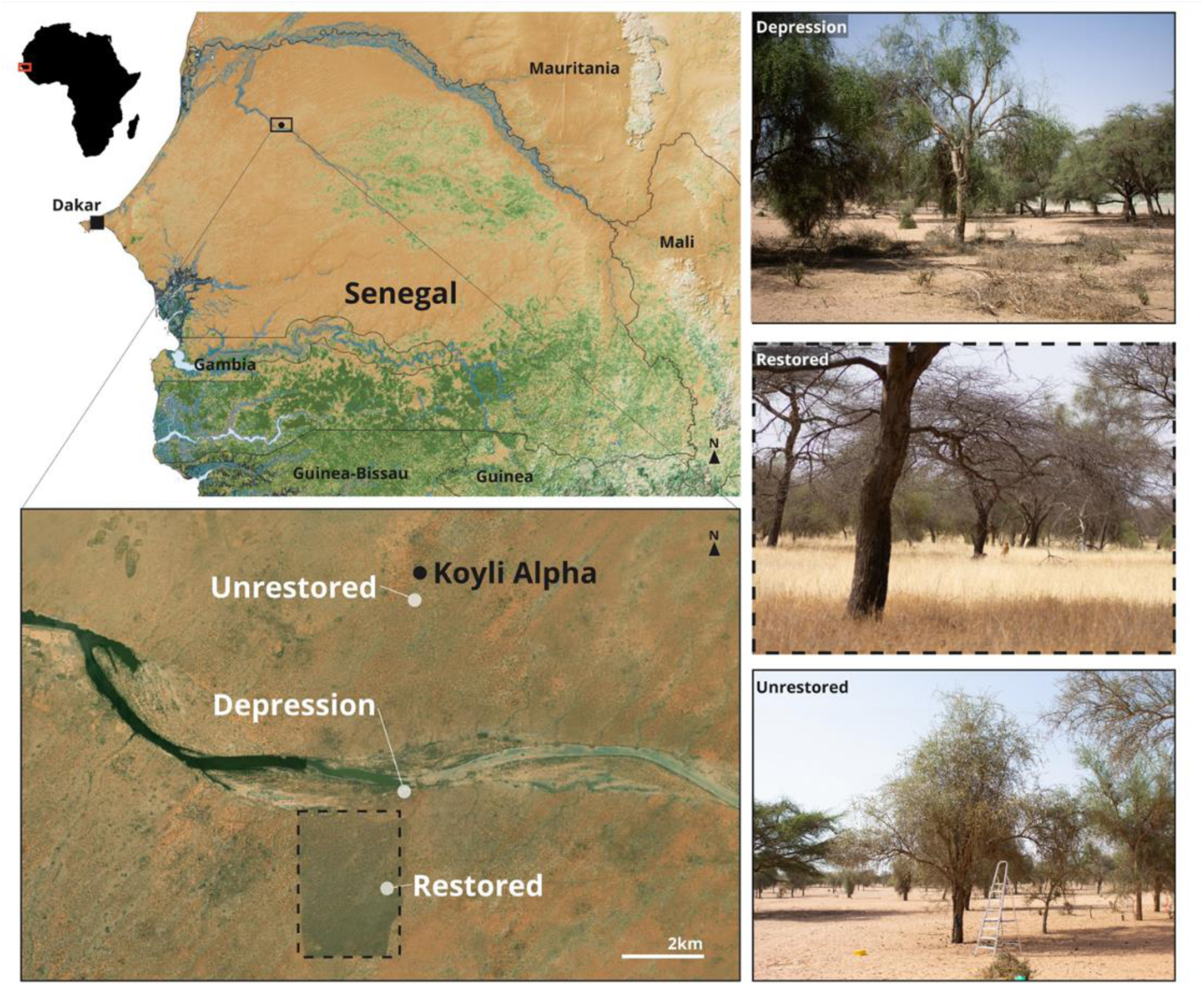
Location of the study area and sampling sites. The map was built under QGIS 3.28.3, using elevation SRTM data at 90m spatial resolution from the CGIAR-CSI database (Jarvis et al., 2008), hydrographic data from (Senegal PCS200, on Africa GeoPortal) and vegetation data from Hansen et al. (2013). Satellite photograph of Koyli Alpha taken in 2019 (Bing Maps), 15.666964°N, - 15.524456°E. Photographs of the field sites were taken during the mid-dry season (February 2023). © N. Medina-Serrano.

### (2) Study species

*Balanites aegyptiaca* (Zygophyllaceae) is a drought-resistant tree species widely distributed in Sahelian ecosystems. The species can be found in various topographic conditions, including hilltops, slopes, and depressions, and can tolerate annual precipitation ranging from 200 to 900 mm, with an optimal rainfall of 400 mm (Dendoncker et al., 2020). Its evergreen foliage and ability to withstand aridity make it a vital component of the region’s ecosystems. Beyond its ecological significance, the tree holds socioeconomic importance, providing construction wood known for its insect-resistant properties, edible fruits sold in local markets, and fodder for goats. This last use is particularly important in the dry season, when the lopped branches of *B. aegyptiaca* are among the few common sources of fodder. Different parts of this tree (fruits, leaves, bark) are also used in traditional medicine (Chothani & Vaghasiya, 2011). *Balanites aegyptiaca* typically grows to an average height of 5.7 m (up to 8-9 m) and features a spherical crown with supple, drooping branches adorned with straight spines measuring 8 to 10 cm in length. Its flowers, grouped in small axillary racemes, are small, hermaphrodite and actinomorphic, offering open access to floral resources (nectar and pollen) by diverse insects. Flowering occurs throughout the year, in both wet and dry seasons. These characteristics underscore the tree’s adaptability and its multifaceted contributions to ecosystem functioning and services provided to humans (Le Houérou, 1989; Arbonnier, 2000; Ndoye et al., 2004; Gebrekirstos et al., 2008; Dendoncker 2020; Dendoncker et al., 2020).

### (3) Study design

To capture seasonal variation in insect diversity and activity patterns, we sampled in three different seasons when the trees were in flower: June 2022 (end of the dry season), August 2022 (rainy season), and February 2023 (middle of the dry season).

Two sampling methods were employed simultaneously: active capture of flying insects using hand nets and passive capture of insects in coloured pan traps. These two complementary methods are frequently used in combination to study species assemblages of flower visitors (Westphal et al., 2008). Netting of insects was conducted at each site around five flowering individuals of the target tree species. Sampling was conducted for a duration of fifteen minutes per tree at heights ranging from two to five meters. In each of the three seasons, this procedure was repeated over three consecutive days, with the order of sampling rotated between sites. To minimise potential bias caused by variations in insect activity throughout the day, the order of sampling was alternated between sites, and between trees within sites, across consecutive days (in a range between 08h00 and 17h00), as it was not possible to sample all trees simultaneously at the same hour. Five trees were sampled per site and per season, for a total of 45 trees.

Pan trapping used round plastic pans (diameter 15 cm) sprayed with UV fluorescent paint (white, yellow, or blue, one of each colour) and containing water and a drop of unscented detergent. The three coloured pans were placed near inflorescences at mid-canopy level at the beginning of each sampling session on three of the five individual trees sampled by hand netting in each site and season to attract and capture flower-visiting insects (Westphal et al., 2008), for a total of 27 trees. The coloured pans were placed using a 2.5 m stepladder in order to attach each pan to branches that were as close as possible to flowers. Pan traps were placed on different branches distributed over the tree crown and at a range of heights similar to that sampled by netting. Use of traps of three different colours helped to limit bias in the identity of the insects collected (Vrdoljak & Samways, 2012). Sampling design is schematised in Figure 2. After 48 hours, we collected the insects from all pan traps and stored them in 80% ethanol. Insects collected with nets were also stored in 80% ethanol at the end of each session. The stored insects were then kept at room temperature and away from direct sunlight until reaching the laboratory to prevent DNA degradation and to maintain the specimens for subsequent observations and identifications.

**Figure 2.**
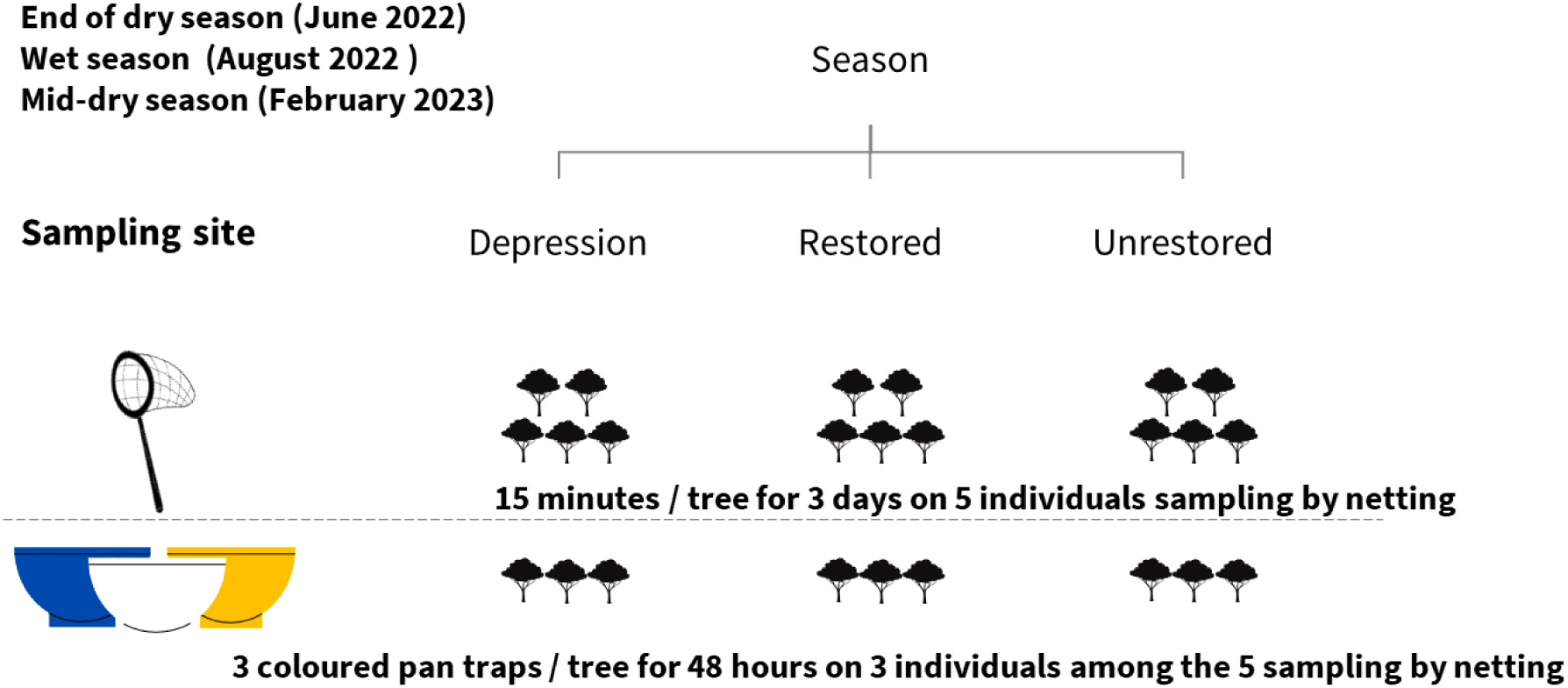
Sampling design for collection of insects visiting flowers of *Balanites aegyptiaca*. For hand net sampling, the order of sampling in the different sites was rotated each day to avoid a confounding effect of time of collection.

The flowering trees sampled were often not the same individuals in the three seasons. For each tree in each sampling period, we estimated the abundance of open flowers on the tree (percentage of the tree’s canopy that bore flowers, semi-quantitative scale that varied from 5% to 85%). For each sampled tree, we also noted the 10 other trees (height ≥ 4 m) closest to it, recording their species identity and the distance of each from the focal tree (Figure S3). This allowed estimation of tree species diversity and density in the neighborhood of each sampled tree.

Owing to logistical constraints, we were able to sample only three sites (one “Depression”, one “Restored” and one “Unrestored”), each sampled during the three seasonal periods. We recognise the limitation imposed by the lack of replication of sites, and attempted to mitigate it by sampling individual *Balanites* trees scattered in each site (distances between trees in the same site ranging from 1 to 15 m) and treating them as replicates. The sampling effort was determined based on the availability of individuals with abundant flowers (≥ 10% of the tree’s canopy) at the same time in a given site.

### (4) Insect taxonomic and functional diversity

#### (a) Insect porcessing and taxonomic identification

Determination of morphospecies involved several steps. The collected insects, stored in 80% ethanol, were first dried and observed and, if necessary, photographed using a Leica stereomicroscope equipped with a Leica MC170 HD camera connected to a computer. After observation, the insects were again placed in ethanol. All determinations were made at the family level or lower taxonomic levels using Aberlenc keys (2021). For Hymenoptera, we used online keys in Atlas Hymenoptera (http://www.atlashymenoptera.net/default.aspx). For Diptera, we also consulted Kirk-Spriggs & Sinclair (2017-2021), with additional verifications for Chloropidae and Milichiidae performed by J. W. Ismay (Oxford University Museum of Natural History) at the end of the 2022 field campaign. Furthermore, Coleoptera were compared with those from the reference collection of the terrestrial invertebrate zoology laboratory at IFAN Ch. A. Diop on the UCAD campus in Dakar, Senegal. For Lepidoptera, we tallied the numbers of individuals captured. Almost all were small moths. We lacked the time to spread and mount them, and it was not possible to distinguish morphospecies with ethanol-preserved specimens. We thus excluded Lepidoptera from our analyses of morphospecies diversity.

Within each family, morphospecies were distinguished and assigned individual codes including the first six letters of the insect family name followed by a unique number ID. The number of specimens per morphospecies was meticulously recorded, and one individual of each morphospecies was photographed for the purpose of documentation. As new morphospecies were recognised based on differences in size, colour, or other traits, the morphospecies list was updated accordingly. All samples are preserved in the personal collection of the first author at the Center for Functional and Evolutionary Ecology (CEFE) in Montpellier, France.

#### (b) Functional group classification

We characterised insect functional diversity by assigning morphospecies to broad functional groups that, in many cases, correspond to family-level classifications. Assignments of morphospecies to functional groups were based on both the literature and our own field observations. These taxonomic and functional classifications allowed a closer look at the ecological roles and interactions of the insect community within the study area.

Functional groups were defined based on the known ecology of each family (see Table 1). Specific classifications, such as “Likely important pollinators”, emphasise the heavy reliance of these insects on floral resources (bees and flies are categorised separately, with bees relying exclusively on floral resources) and their status as pollen carriers reported in the literature and confirmed through our observations. The capacity to transport pollen was inferred from family-level or genus-level traits and morphological similarities to taxa with established pollination roles. Families in which species vary widely in their ecology are classified as “diverse.” Our classification has obvious limitations. For example, Tachinidae (Diptera) are parasitoids but are frequently mentioned as flower visitors and potential pollinators, justifying their classification as “likely important pollinators” along with other dipteran families whose adults are heavily dependent on floral resources. Similarly, species of families classed as predatory or as phytophagous are also known to contribute to pollination (e.g., predatory wasps). The group “parasitoids” encompasses insects whose hosts include a great diversity of arthropod species and life-cycle stages. Despite these limitations, we believe our classification provides a general idea of the guilds represented by the different groups visiting the flowers of *B. aegyptiaca*. We use our functional-group classification only to support discussion, and not statistical analyses.

**Table 1.**
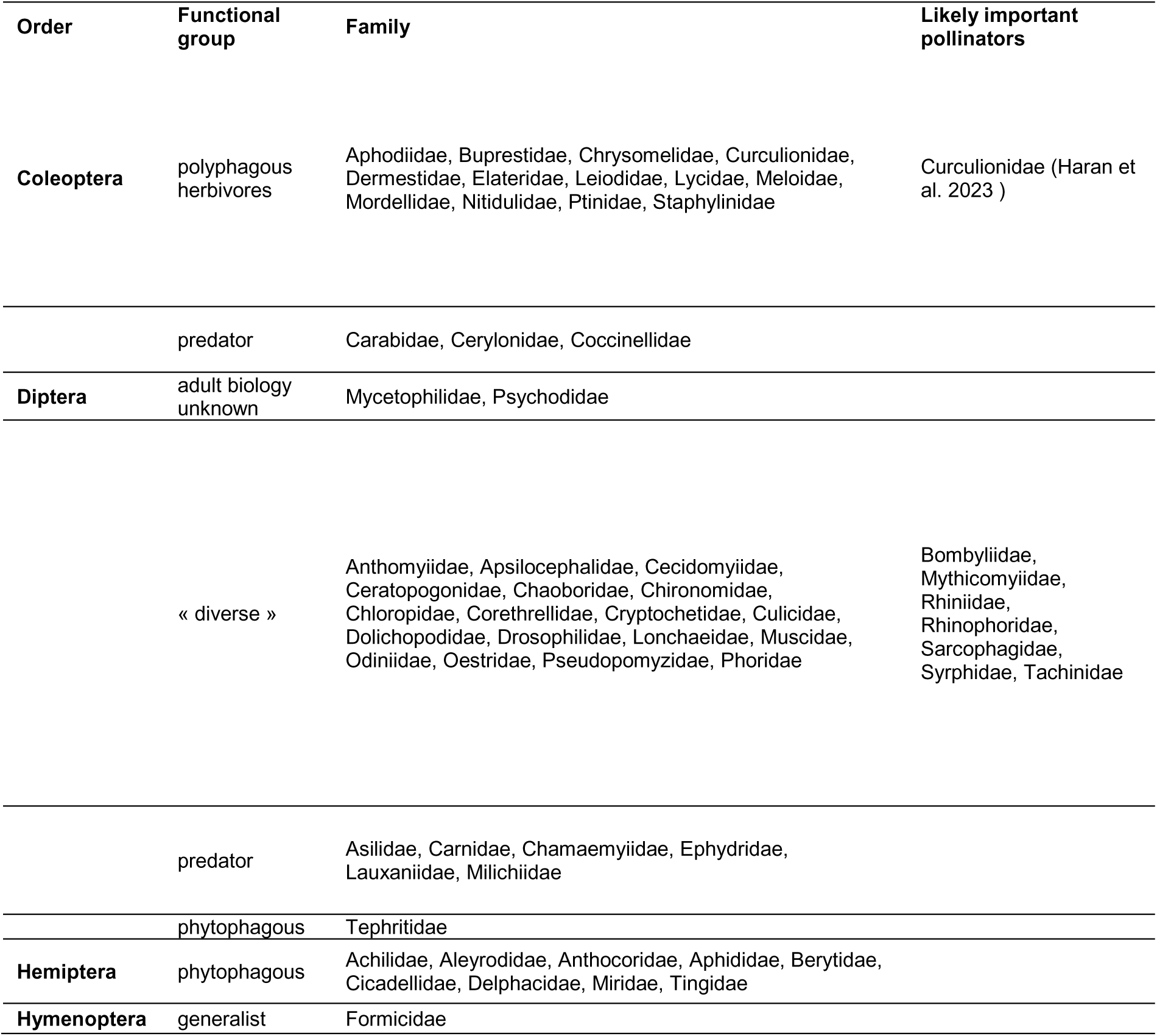
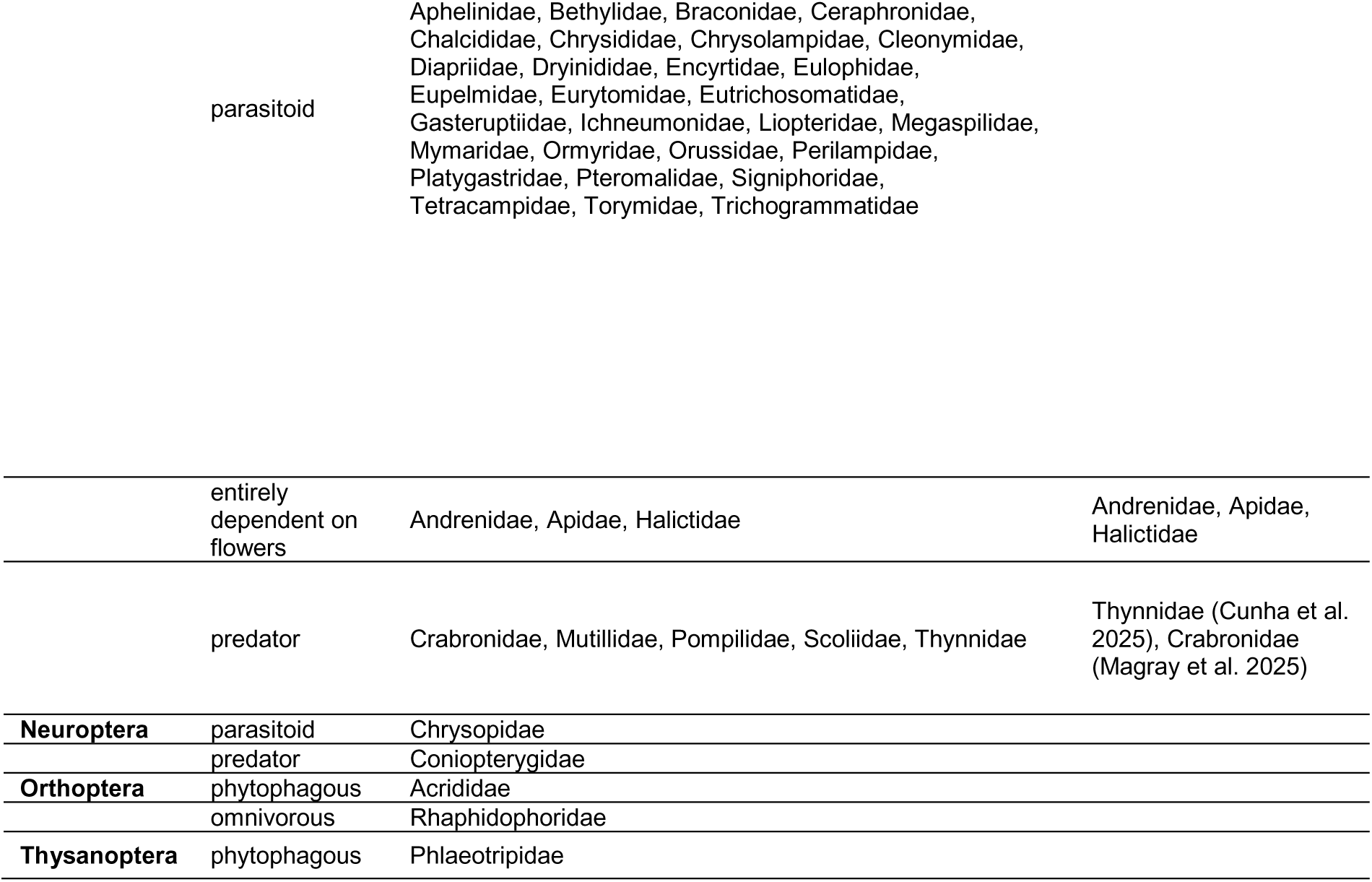
Flower-visiting insects captured (pan-trap and hand net samples), grouped by orders, broad functional groups, and families. Biological information on families taken from Aberlenc (2021), Kirk-Spriggs & Muller (2017-2021), and Mawdsley et al. (2016). Classification of “likely important pollinators” was based on the information provided in the column “Importance in pollination” in Tables 5, 6 and 8 in Medina-Serrano et al. (2025).

### (5) Analysis of global patterns of insect abundance, composition and diversity

#### (a) Data description and analysis settings

Analyses were conducted in R version 4.5.0 (R Core Team, 2025) using RStudio (Version 2025.05.0+496 ‘Mariposa Orchid’; RStudio Team, 2025).

Species abundances were aggregated to represent the total abundance for each order. Then, species abundance distributions were used to estimate true species richness and diversity. Species-abundance distributions in our data (hand net and pan-trap captures combined) followed a log-normal distribution, characterised by many rare species and a few very abundant ones (see Figure S4). When dealing with small sample sizes, caution is needed in extrapolating the overall shape of a species abundance distribution from limited data. The negative binomial distribution is considered the best fit when there is a large proportion of rare species (Chao et al., 2014b; Callaghan et al., 2023). We thus computed our models of abundance using this distribution.

#### (b) How complete was our sampling? Diversity and sampling completeness

Sampling usually does not exhaustively cover the true species richness of organisms (Schmitz & Rahmann, 2025). We assessed the completeness of our coverage of species richness of flower-visiting insects using rarefaction and cumulative curves. The Chao1 richness was computed to estimate true species richness, based on the observed richness and abundance distributions, with species ranked in order of abundance from the rarest to the most abundant for each season separately (Chao et al. 2014a).

Cumulative curves are shown for both methods combined (hand net and pan-trap) and for each of the three seasons: end-dry season (June), rainy season (August) and mid-dry season (February), indicating the total number of individuals sampled and the number of species observed for each season. We then compared observed species richness and the Chao1 richness estimator for each season for the two methods combined.

Subsequently, to quantify the cumulative richness per site, we pooled data for all trees in each site (“Restored”, “Unrestored”, and “Depression”) for both sampling methods combined, indicating the total number of individuals sampled and the number of species observed for each site. We then compared observed species richness and the Chao1 richness estimator for each site for the two methods combined.

#### (c) Standardisation of sampling effort

To comprehensively describe the dataset, we first combined data from both sampling methods in all seasons and for all sites in global analyses of the abundance, diversity, and composition of insects visiting the flowers of B. aegyptiaca. To account for differences in sampling effort among sites, seasons, and methods, abundance and diversity data were standardised prior to analysis. Diversity indices were standardised by sample coverage using the *iNEXT* package (Chao et al., 2014a), to calculate three indicators of diversity, species richness (Hill number = 0), Shannon index (Hill number = 1) and the Simpson index (Hill number = 2), utilising the iNEXT R package (Hsieh et al., 2016) under a common level of sampling completeness using the function estimateD. Data on insect abundance and diversity from each tree taken over three days were aggregated for analysis. Treating each tree as a replicate, we used Generalised Linear models (GLMs) and Linear models (LMs) to examine the influence of season and site. For each site and season, insect abundance was first calculated at the tree level. To allow direct comparison between hand-net and pan-trap sampling, abundances from both methods were standardised to represent the mean catch per tree (i.e., total abundance divided by five for the hand-net sampling and by three for the pan-trap sampling). This ensured that abundance estimates for both methods corresponded to an equivalent tree-level sampling unit.

#### (d) Effects of vegetation structure and floral resources

Differences in mean distance and flowering canopy among sites were analysed with Kruskal–Wallis tests. We then fitted separate GLMs by season and site to test the effects of local tree density (mean distance to the ten closest neighbouring trees) and canopy flowering percentage. These models were used to evaluate whether the habitat variables explained variation in mean insect abundance (see Table S1 for details). These variables were not included in the subsequent model combining season and site (see Section (e)).

#### (e) Effects of season and site

Abundance of insects was modelled using generalised linear models (GLMs). The GLMs were performed with the glm.nb function with the MASS R package, which fits a negative binomial regression model (using log = ln ; Ripley & Venables, 2009), whereas species richness and Simpson diversity were analysed with linear models (LMs) assuming normal residuals. The linear regression models were performed with the lm function using the log(x+1) transformed data before analysis to normalise variance and reduce heteroscedasticity. Models included season, site, and their interaction as fixed effects.

Residuals of models were verified to confirm the appropriateness of the assumptions using the R package DHARMa (Brooks et al., 2017; Hartig & Lohse, 2020). When significant differences were found across seasons or across sites, we conducted pairwise comparisons (comparing each pair of seasons or of sites) using the *pairs* function from the emmeans R package (Lenth, 2023). Owing to the limited number of observations, models were made as simple as possible, examining effects of only season and site and their interaction and combining data from the two sampling methods. To assess the effects of site, season, and their interaction on insect abundance, we fitted a Generalised Linear Model (GLM) with a negative binomial error distribution using the MASS package (Ripley & Venables, 2009). The model addresses potential overdispersion in the count data by employing the negative binomial distribution, which is more appropriate than the Poisson distribution for this type of data (Baldridge et al., 2016). The response variable was the number of insects observed per tree, with site (Depression, Unrestored, Restored) and season (end-dry, rainy, mid-dry) included as fixed effects. The model therefore tested the independent effects of site and season as well as their interaction.

Model fit and the proportion of variance explained were quantified using the MuMIn package (https://cran.r-project.org/web/packages/MuMIn/index.html), which provides both marginal R² and conditional R². Significance of the predictors was evaluated through a Type III analysis of deviance with chi-square tests using the car package (Fox et al. 2019), which allowed us to assess the relative contributions of site, season, and their interaction to explaining variation in abundance.

To evaluate whether model assumptions were met, we performed residual diagnostics with the DHARMa package (Brooks et al., 2017; Hartig 2024). Simulated residuals were generated and plotted, providing a robust check for potential issues such as overdispersion, zero inflation, or non-uniformity in residuals.

Following model validation, we examined post-hoc comparisons using estimated marginal means with the emmeans package. We first compared average abundances across sites (averaged over seasons) and across seasons (averaged over sites) followed by pairwise comparison. To explore the interaction, we performed pairwise comparisons of sites within each season (*emmeans (model, pairwise ∼ site | season*)) and of seasons within each site *emmeans* (*model, pairwise ∼ season | site*), applying Tukey’s adjustment to control for multiple testing. Taken together, this framework allowed us to test for both main and interactive effects of site and season on insect abundance and evaluate effect sizes on the response scale, while accounting for overdispersed count data and the risk of inflated Type I errors due to multiple comparisons.

For the diversity indices, we used two linear regression models with log(x+1) transformation of the data to examine the effects of season and site and their interaction on species richness (Hill number = 0), Shannon diversity index (Hill number = 1) and Simpson diversity index (Hill number = 2). This transformation helped manage skewness and outliers in the data. The models tested whether richness or diversity varied independently with site and season, as well as whether seasonal differences depended on site type.

Model diagnostics were performed by plotting simulated residuals using the DHARMa package to verify that assumptions of linear modelling were met, including homoscedasticity and normality of residuals. The amount of variation explained by the models was quantified using marginal and conditional R² values in the MuMIn package. The significance of main effects and of their interaction was assessed through Type III analysis of variance (ANOVA) with chi-square tests from the car package. Post-hoc comparisons were then carried out with the emmeans package. Estimated marginal means (EMMs) were computed for each factor level, and pairwise comparisons were performed both across sites within each season and across seasons within each site type, applying Tukey’s adjustment for multiple comparisons.

#### (f) Seasonal contrasts

##### Seasonal patterns in species composition of insect assemblages

To illustrate overall patterns of seasonal variation in the morphospecies composition of the captured insects, we tabulated the number of morphospecies unique to each season and the number shared by each combination of seasons, constructing a Venn diagram. Analyses were carried out using the function *vegdist* (vegan R package; Oksanen et al. 2025).

##### Seasonal patterns in specific taxonomic groups

We tested the effects of season on the abundance of the main flower-visiting insect orders and families to obtain a more detailed picture of seasonal patterns. These analyses were performed using a negative binomial GLM, with order- or family-level abundance as response variable, and season (as factor) as the explanatory variable, following the methods cited above. Seasonal effects were tested for the orders Hymenoptera, Diptera, Hemiptera, and Coleoptera. Family-level analyses were possible for only three families (all Hymenoptera): Braconidae, Halictidae and Formicidae.

## Results

### (1) Community of insects visiting flowers of *Balanites aegyptiaca*

We found an unexpectedly high diversity of insects at the flowers of this tree species in this extremely harsh environment. Across three sites and seasons (five trees per site per season), we captured 2907 individuals from a total of at least 371 morphospecies from 10 insect orders (Figure 3).

**Figure 3.**
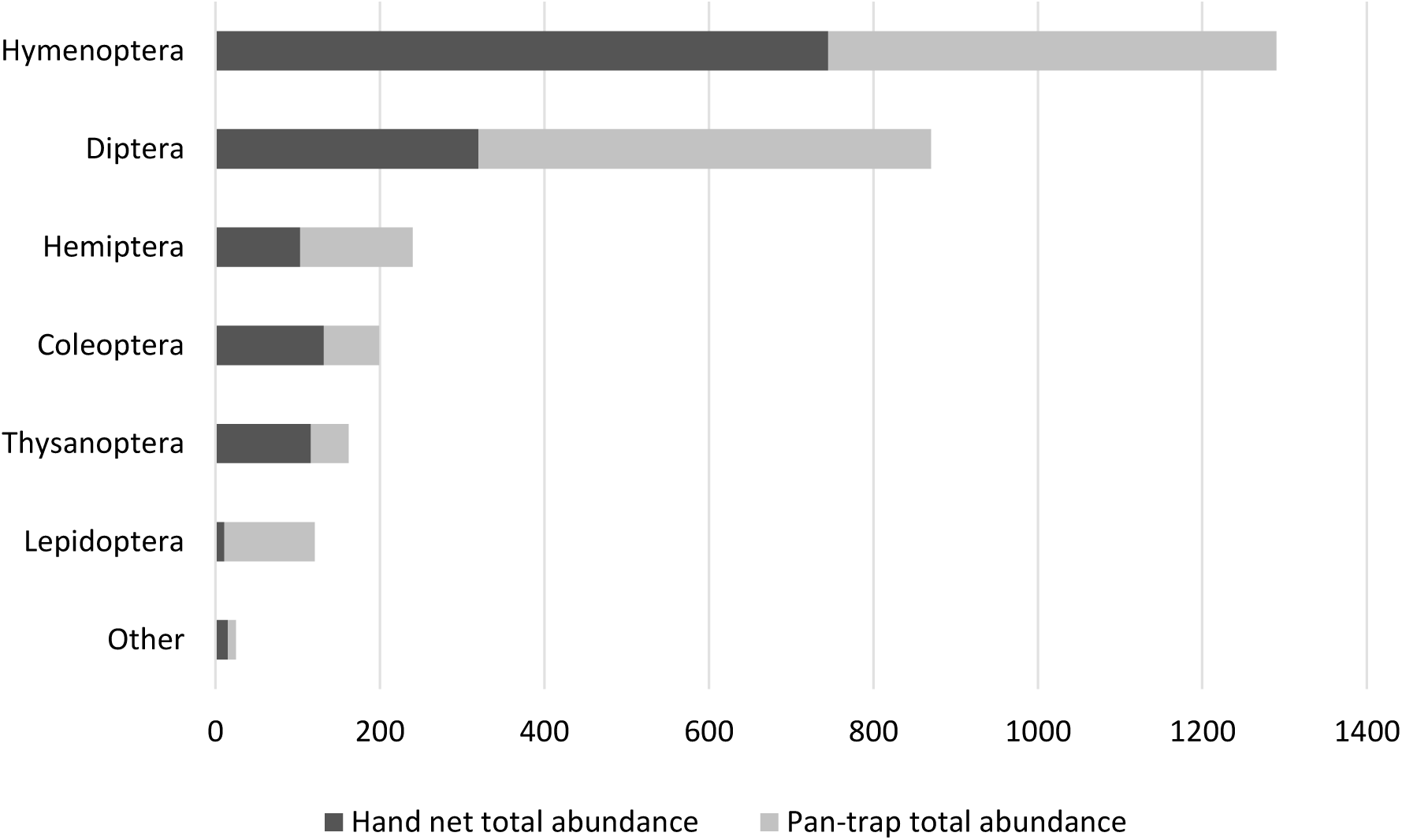
Barplot of the total abundance of insects of different orders captured at flowers of *Balanites aegyptiaca.* “Other” includes Ephemeroptera, Neuroptera, Orthoptera and Psocoptera. The two methods (hand net and pan-trap) combined for a total of 2907 individuals of at least 371 morphospecies (morphospecies not distinguished for Lepidoptera). A total of 1442 insects were collected by hand net sampling, and a total of 1465 insects were collected by pan-trap sampling.

The most abundant and diverse orders were Hymenoptera (1290 individuals, 180 morphospecies) and Diptera (870 individuals, 101 morphospecies), followed by Hemiptera (240 individuals, 30 morphospecies), Coleoptera (199 individuals, 39 morphospecies), Thysanoptera (162 individuals, 11 morphospecies), and Lepidoptera (121 individuals, morphospecies not distinguished). Four other orders (Ephemeroptera, Neuroptera, Orthoptera and Psocoptera) contributed a combined total of 25 individuals and 10 morphospecies (Figure 3). Table 1 presents the orders and families of flower-visiting insects encountered in the study, and the functional categories we used to describe them.

There were some notable differences in the assemblages captured by the two methods (Figure 3). First, Lepidoptera, almost all of which were small moths, comprised only 0.8% of the hand net samples but 7.5% of pan-trap samples. This difference may reflect the nocturnal habits of most species captured. Netting was done only during the day, whereas pan traps sampled insects over 48 hours. Lepidoptera were captured primarily at the end of the dry season, with 85 individuals recorded in June, compared to 35 individuals during the mid-dry season and only three individuals during the rainy season. Secondly, the two most abundant orders, Hymenoptera and Diptera, were equally abundant in pan-trap samples but Hymenoptera were two times more abundant than Diptera in hand net samples.

As stated in the Methods section, Lepidoptera were not further analysed. Insects of all nine other orders were captured by both sampling methods. The two sampling methods contributed a roughly equal number of individuals to the total sample (pan traps: 1465; hand net: 1442). Note, however, that at each season and site, five trees were sampled by netting, while a subset of three of these trees was sampled using pan traps. The number of insects captured per tree was thus about 1.7 times higher for pan traps than for hand nets.

### (2) Effects of season and site on abundance and diversity flower-visiting insects on Balanites aegyptiaca

#### (a) Comparison of observed and estimated species richness across seasons

Observed species richness (both capture methods combined) tended to be lower at the end of the dry season (104 species), followed by the rainy season (180 species) and highest during the mid-dry season (207 species) (Figure 4).

**Figure 4.**
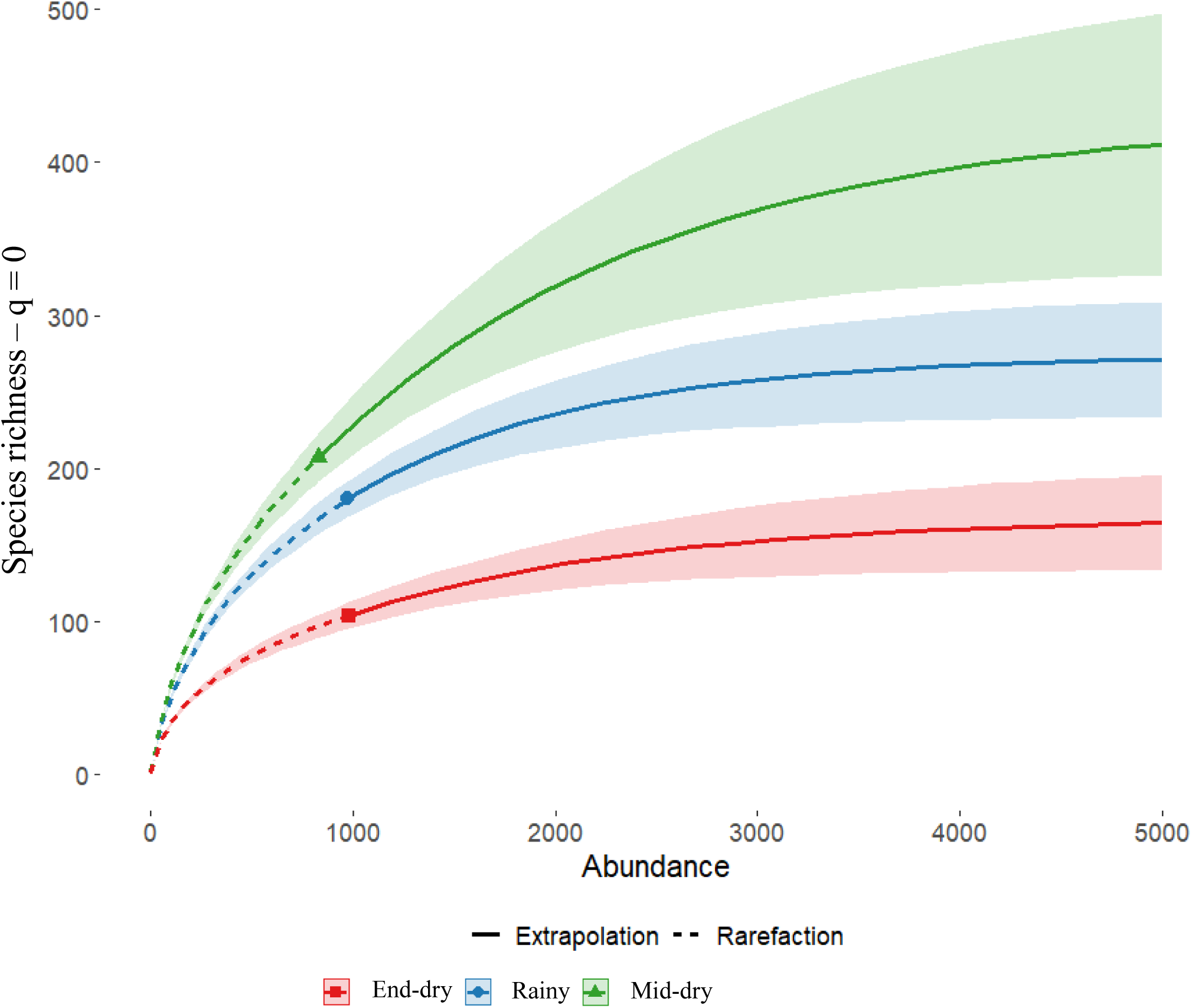
Accumulation curves (results for both capture methods combined) showing seasonal variation in richness of insect morphospecies at flowers of *Balanites aegyptiaca*. Red represents the end-dry season (June); blue represents the rainy season (August); and green represents the mid-dry season (February).

Estimated richness (ChaoRichness) was also lowest at the end of the dry season (168 species). However, in contrast to the relatively similar observed richness between the rainy and mid-dry seasons, estimated richness was substantially higher during the mid-dry season (431 species) than during the rainy season (274 species) (Figure 4).

#### (b) Comparison of observed and estimated species richness across sites

Observed species richness (both capture methods combined) tended to be lower in the Unrestored site (171 species) and similar in the Restored and Depression sites (201 and 203 species, respectively). Estimated richness (ChaoRichness) was also lowest in the Unrestored site (273 species). However, in contrast to their similar observed richness, estimated richness was much higher in the Depression site (419 species) than in the Restored site (317 species) (Figure 5).

**Figure 5.**
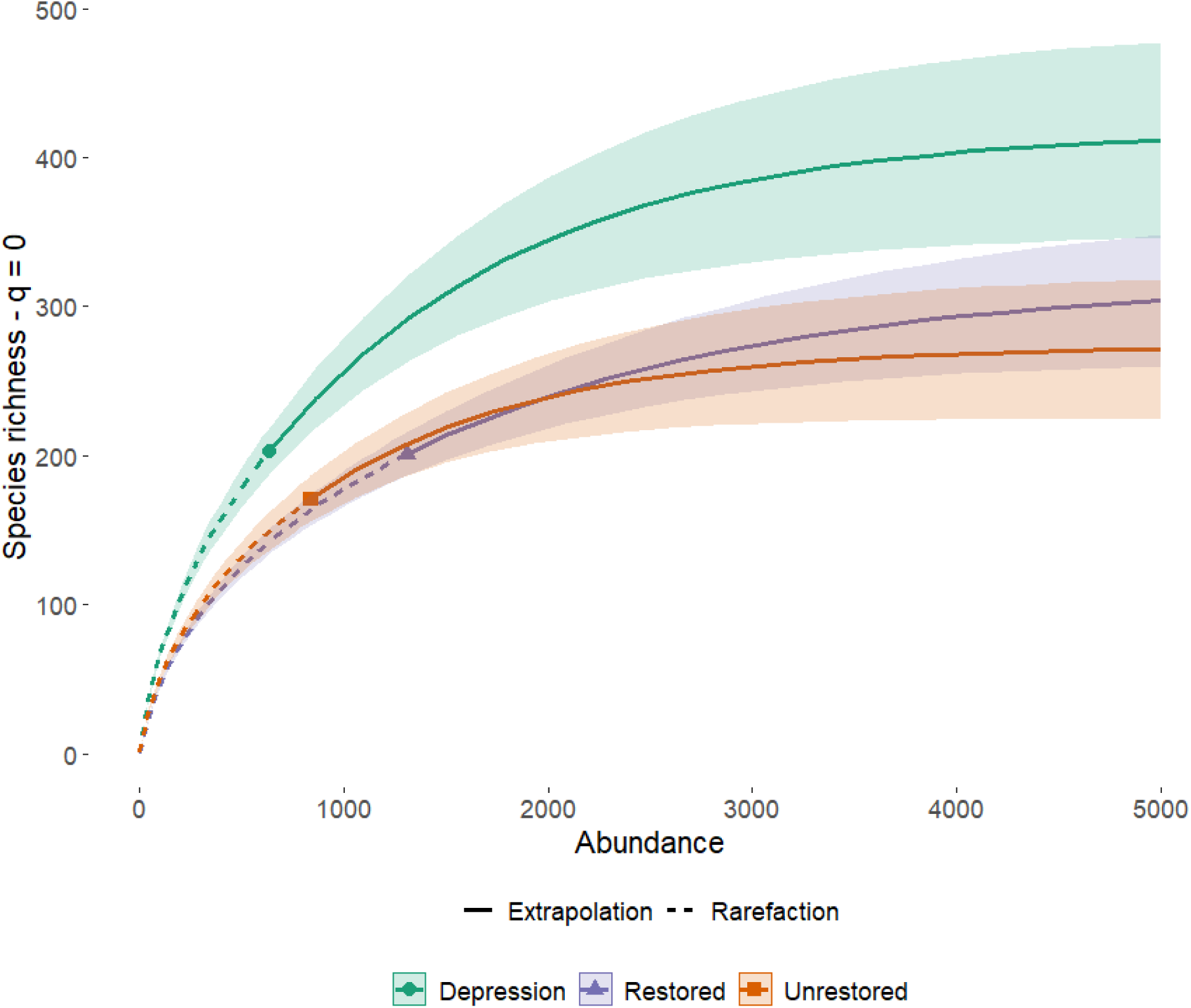
Accumulation curves showing variation among sites in morphospecies richness on *Balanites aegyptiaca*. In green the Depression site, in purple the Restored site, and in orange the Unrestored site.

#### (c) Effects of vegetation structure and floral resources

Mean distance to neighbouring trees differed significantly among sites, whereas canopy flowering percentage did not (Kruskal–Wallis tests). Mean distance was significantly lower at the Depression site than at the two other sites (Figure S5). Insect abundance per tree was positively correlated with mean distance to the ten closest neighbouring trees when analysed across seasons, but not across sites (GLM models). In analysis of the seasonal effect, insect abundance per tree increased significantly with greater mean distance to neighbouring trees (χ² = 5.81, df = 1, p = 0.016). In contrast, canopy flowering percentage had no significant effect on insect abundance per tree in either the seasonal or site-specific models (see Table S1 for details). Consequently, these two habitat-related variables were not included in the model combining season and site.

#### (d) Effects of season and site on variation in abundance and diversity metrics

The negative binomial generalised linear model (GLM) of insect abundance per tree, including site, season and their interaction, explained a moderate proportion of the variance (R² ≈ 0.34). Effects were tested using likelihood-ratio χ² statistics (analysis of deviance). The analysis revealed a significant main effect of site on insect abundance (LR χ² = 14.01, df = 2, p < 0.001), whereas the effects of season (LR χ² = 5.53, df = 2, p = 0.063) and the site × season interaction (LR χ² = 9.25, df = 4, p = 0.055) did not reach the 0.05 significance level (see Table S1).

Across sites (i.e., averaged over seasons), mean insect abundance per tree was significantly lower in the Depression than in the Restored site (p = 0.0029), with a similar but non-significant trend for the Depression compared to the Unrestored site (p = 0.0719). Mean insect abundance did not differ between the Restored and Unrestored sites (p = 0.62). No overall differences in insect abundance were detected among seasons when averaged across sites (all pairwise comparisons: p ≥ 0.86; see Table S2). Within-season contrasts showed a strong site effect at the end of the dry season (June): mean abundance per tree in the Depression site was significantly lower than in both the Restored (p = 0.0004) and Unrestored sites (p = 0.0104), while no significant difference was found between the Restored and Unrestored sites in that season (p = 0.6104). No significant site differences in insect abundance were detected in the rainy (August) or mid-dry (February) seasons (all p > 0.11; Figure 6a; Table S2).

**Figure 6.**
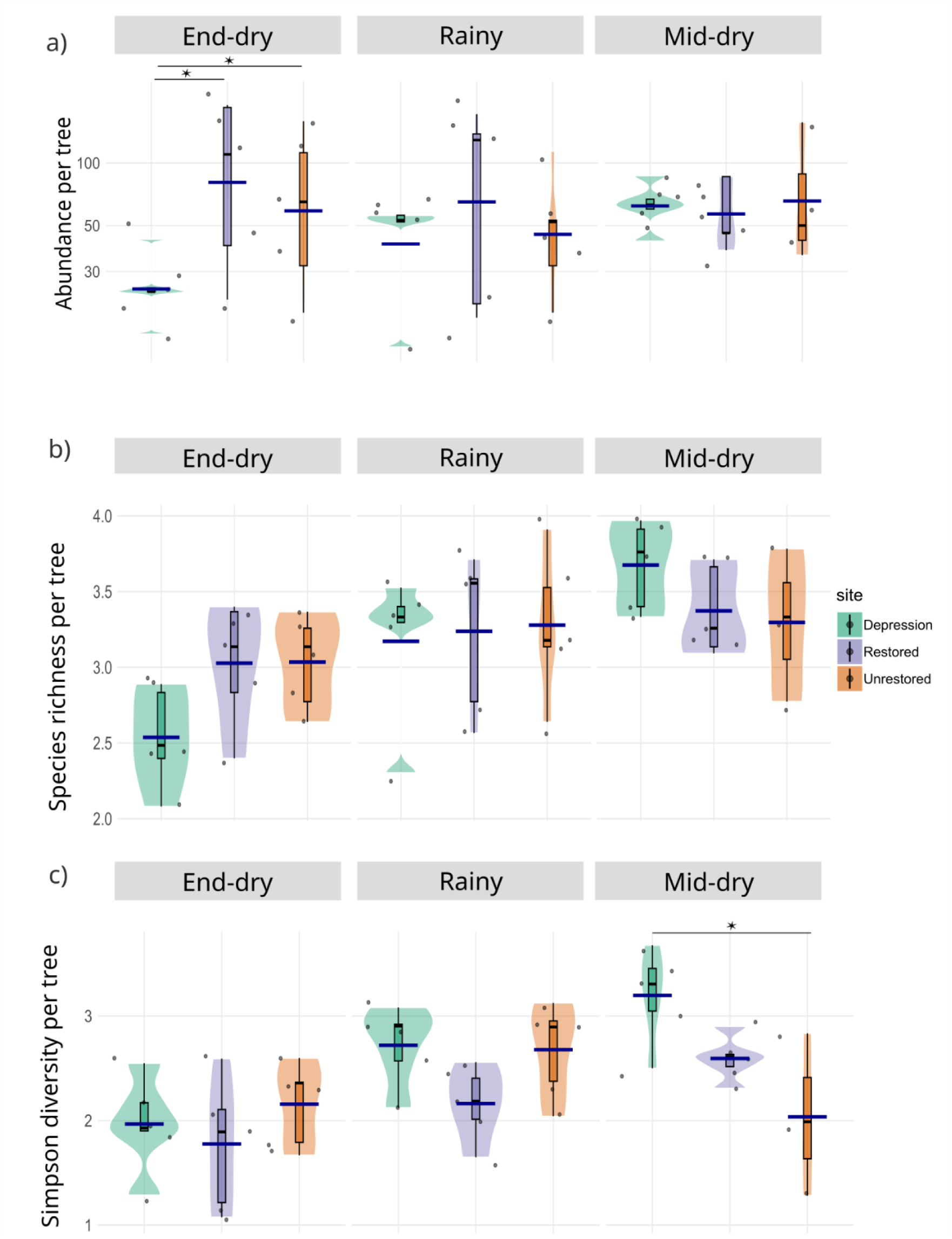
Violin plots of insect assemblages at flowers of Balanites aegyptiaca, showing for each site (a) log 10 (abundance per tree), (b) log1p(observed species richness (q = 0) per tree), and (c) log1p(Simpson diversity index (q = 2) per tree) across seasons: end-dry (June), rainy (August), and mid-dry (February). Each violin illustrates the probability density of values within a season. Boxplots inside each violin show the median and interquartile range, and individual sampling points are overlaid as jittered dots to reduce overlap. Dark blue horizontal bars indicate the mean of each group. Colours indicate site categories (green = Depression, orange = Unrestored, purple = Restored), and facets allow comparisons across seasons. Asterisks indicate significant differences (p < 0.05).

For observed species richness (q = 0), the linear model including site, season, and their interaction explained a moderate proportion of the variation (R² adj. = 0.26; F_(8,34)_ = 2.82, p = 0.016). Season had a strong effect (ANOVA: F = 9.63, p < 0.001), while site and the site × season interaction were not significant (p = 0.105 and p = 0.189, respectively). Pairwise contrasts revealed that at the Depression site, richness in the end-dry season (June) was significantly lower than in the mid-dry season (February, p = 0.0003) and marginally lower than in the rainy season (August, p = 0.051), whereas differences between the rainy season and mid-dry season were not significant (p = 0.144). At the Restored and Unrestored sites, no seasonal contrasts were significant (all p > 0.38). Across sites, no significant differences were detected within any given season (all p > 0.15). These results indicate that variation in species richness was primarily driven by seasonality, but mainly in the Depression site, which showed much greater richness during the rainy and mid-dry season relative to the end of the dry season (Figure 6b; Table S3).

For Simpson diversity (q = 2), the model accounted for 42% of the adjusted variance (R² adj. = 0.42; F_(8,34)_ = 4.77, p < 0.001). Both season (ANOVA: F = 9.06, p < 0.001) and the site × season interaction (F = 2.84, p = 0.039) were significant, whereas site alone was not (p = 0.435). At the Depression site, Simpson diversity was significantly lower in the end-dry season (June) than in the rainy season (August, p = 0.037) and mid-dry season (p = 0.0005), but was not different between the rainy season and the mid-dry season (p = 0.244). At the Restored site, diversity in the end-dry season was significantly lower than in the mid-dry season (p = 0.022) but not significantly different from that in the rainy season (p = 0.393). At the Unrestored site, seasonal differences were not significant (all pairwise comparisons p > 0.15). Within-season contrasts showed that in the mid-dry season, diversity in the Depression site was significantly higher than in the Unrestored site (p = 0.0042), while no significant differences in diversity were observed in any site between the end-dry season and the rainy season (all p > 0.11) (Figure 6c; Table S3).

Shannon diversity (q = 1) generally followed the same trends as Simpson diversity, with significantly higher values during the rainy and mid-dry seasons and lower values at the end of the dry season. Therefore, only the Simpson diversity results are presented here in detail, while Shannon diversity outcomes are provided in the Supporting Information (Figure S6 and Table S3). Overall, these results reveal a consistent and strong seasonal signal across all three diversity indices, with richness and diversity lowest at the end of the dry season (June) and highest during the rainy season (August) and mid-dry season (February), particularly at the Depression site. Site differences were generally weak but became evident in the mid-dry season, when Simpson and Shannon diversity were significantly higher in the Depression site than in the Unrestored site.

### (3) Seasonal patterns in composition of assemblages of flower-visiting insects on Balanites aegyptiaca

#### (a) Seasonal patterns in species of insects assemblages

The composition of insect assemblages varied greatly across seasons. Of 371 morphospecies recorded, only 21 (5.6%) were present in all three seasons. Each season had 46–60% of morphospecies unique to that period. Overlap in morphospecies composition was highest between the rainy and mid-dry seasons, with 20% of species shared. Overlap was lower between the mid-dry and end-dry seasons, with 14.7% of species shared, and between the end-dry and rainy seasons, with 12.8% of species shared (Figure 7).

**Figure 7.**
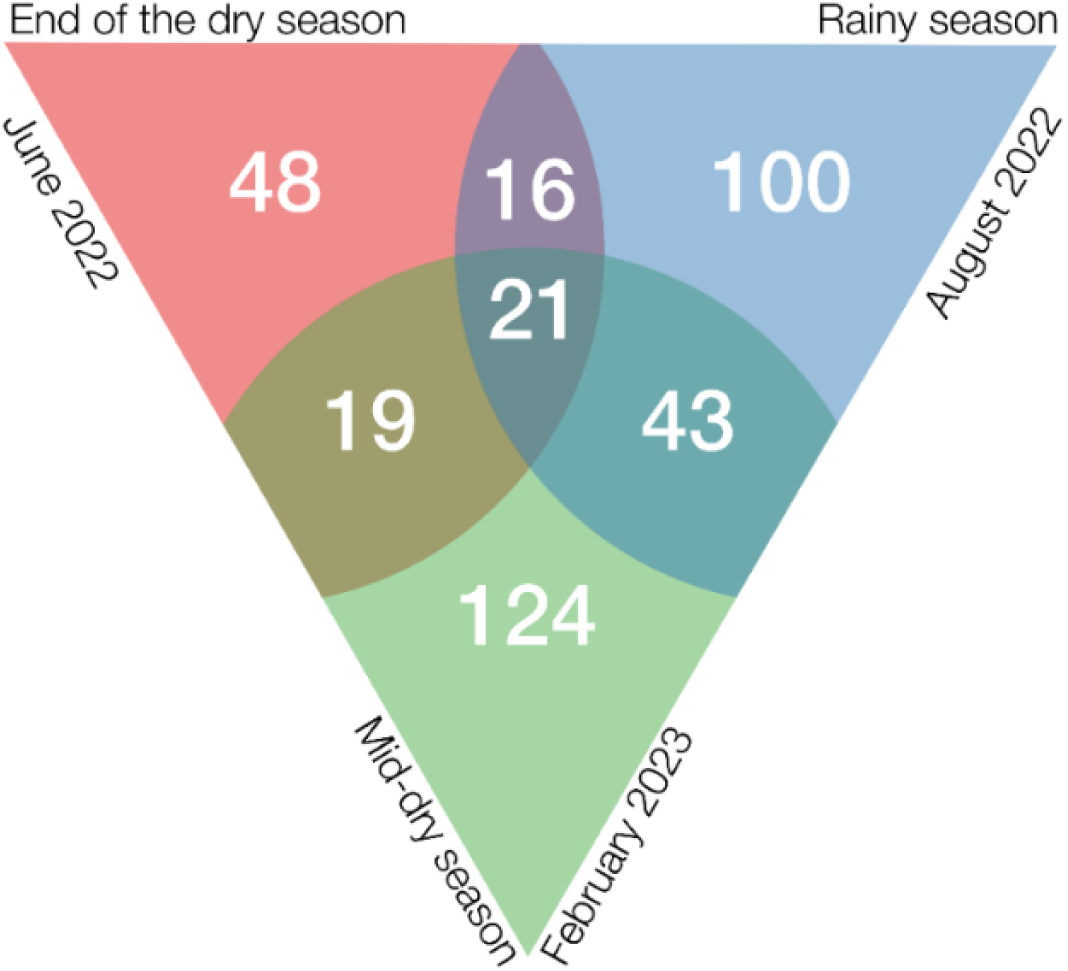
Venn diagram showing great seasonal variation in composition of insect assemblages on *Balanites aegyptiaca*. In pink the end of the dry season, in blue the rainy season and in green the mid-dry season. The numbers give the number of species captured only in a single season, those captured in two seasons, and those captured in all three seasons.

#### (b) Seasonal patterns in selected insect orders

Season affected variation in abundance of different insect orders (Figure 8). Seasonal effects were significant for Hymenoptera (χ² = 15.02, df = 2, p < 0.001), Diptera (χ² = 12.95, df = 2, p = 0.002), and Hemiptera (χ² =15.82, df = 2, p < 0.001) but not for Coleoptera (χ² = 3.80, df = 2, p = 0.15). Pairwise comparisons showed that Hymenoptera were significantly more abundant at the mid-dry season and the end-dry season than in the rainy season (p = 0.036 and p < 0.001, respectively), whereas Diptera and Hemiptera showed the opposite pattern, with particularly low abundance at the end-dry season compared to the rainy season (p < 0.001 for both orders). Additionally, significant differences were found for Diptera and Hemiptera (p = 0.017; p = 0.009 respectively) between the end-dry season and the mid-dry season (see Figure 8 and Table S4). These results suggest that while Hymenoptera, Hemiptera, and Diptera showed more or less strong seasonal shifts, Coleoptera exhibited less seasonal variation.

**Figure 8.**
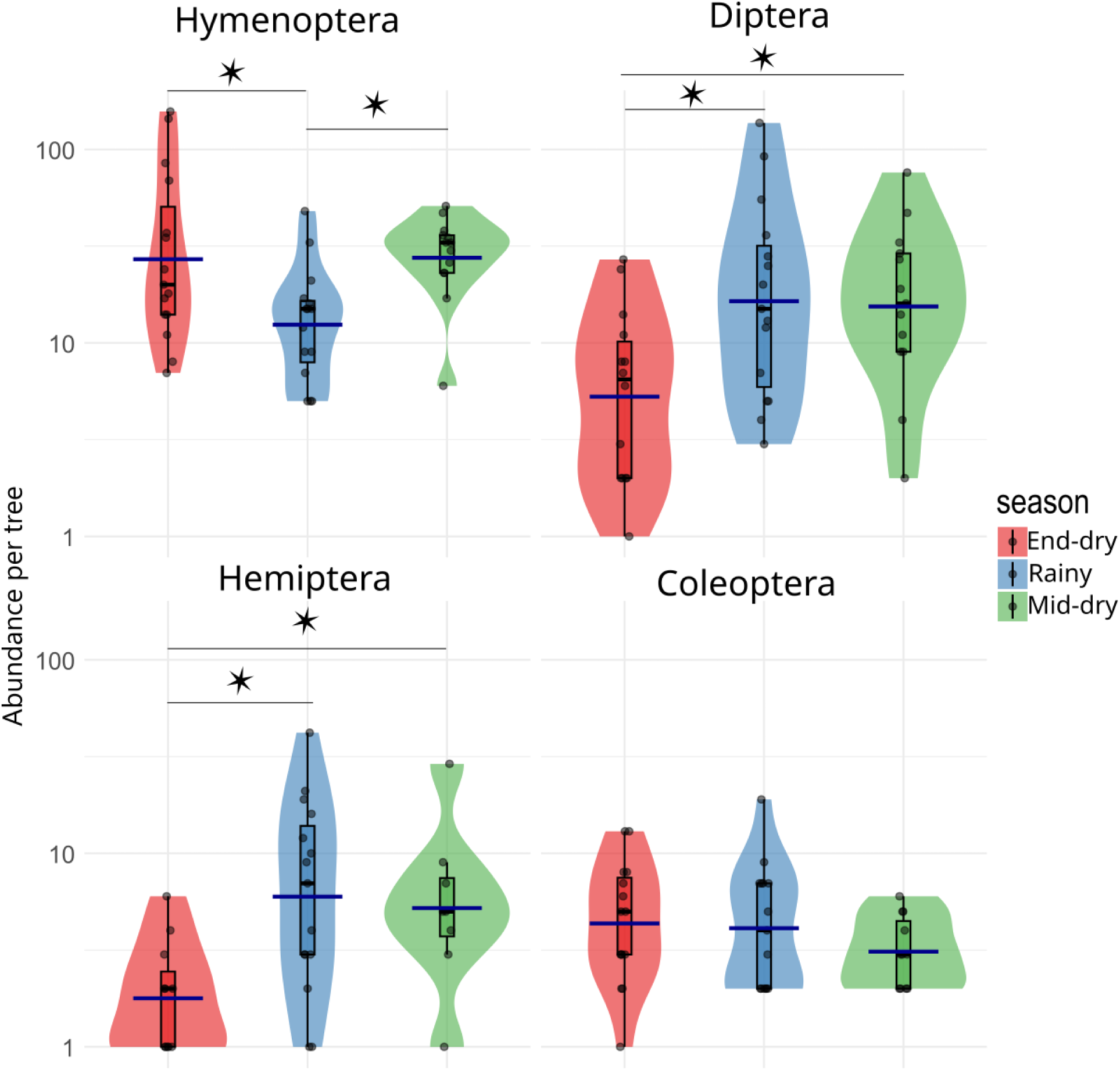
Violin plots showing the distribution in each site of the mean abundance per tree of four orders of insects (Hymenoptera, Diptera, Hemiptera and Coleoptera) visiting flowers of *Balanites aegyptiaca* (combining results of the two sampling methods). In red: end-dry season (June); In blue: rainy season (August); in green: mid-dry season (February). Each violin represents the probability density of abundance values within a season, with individual sampling points overlaid as jittered dots to make overlapping data visible. Boxplots inside each violin show the median and interquartile range. Dark blue bars indicate the mean abundance for each group. See Table S3 for statistical details. Asterisks indicate significant differences (p < 0.05). Count data were analysed using negative binomial generalized linear models with a log link. Data are presented on log 10-scaled axes for visualization purposes.

#### (c) Seasonal pattern in selected families

To provide biological insights, seasonal variation in abundance was examined for four common insect families based on their ecological and biological characteristics (Figure 9). Significant seasonal differences were observed for all four families studied.

**Figure 9.**
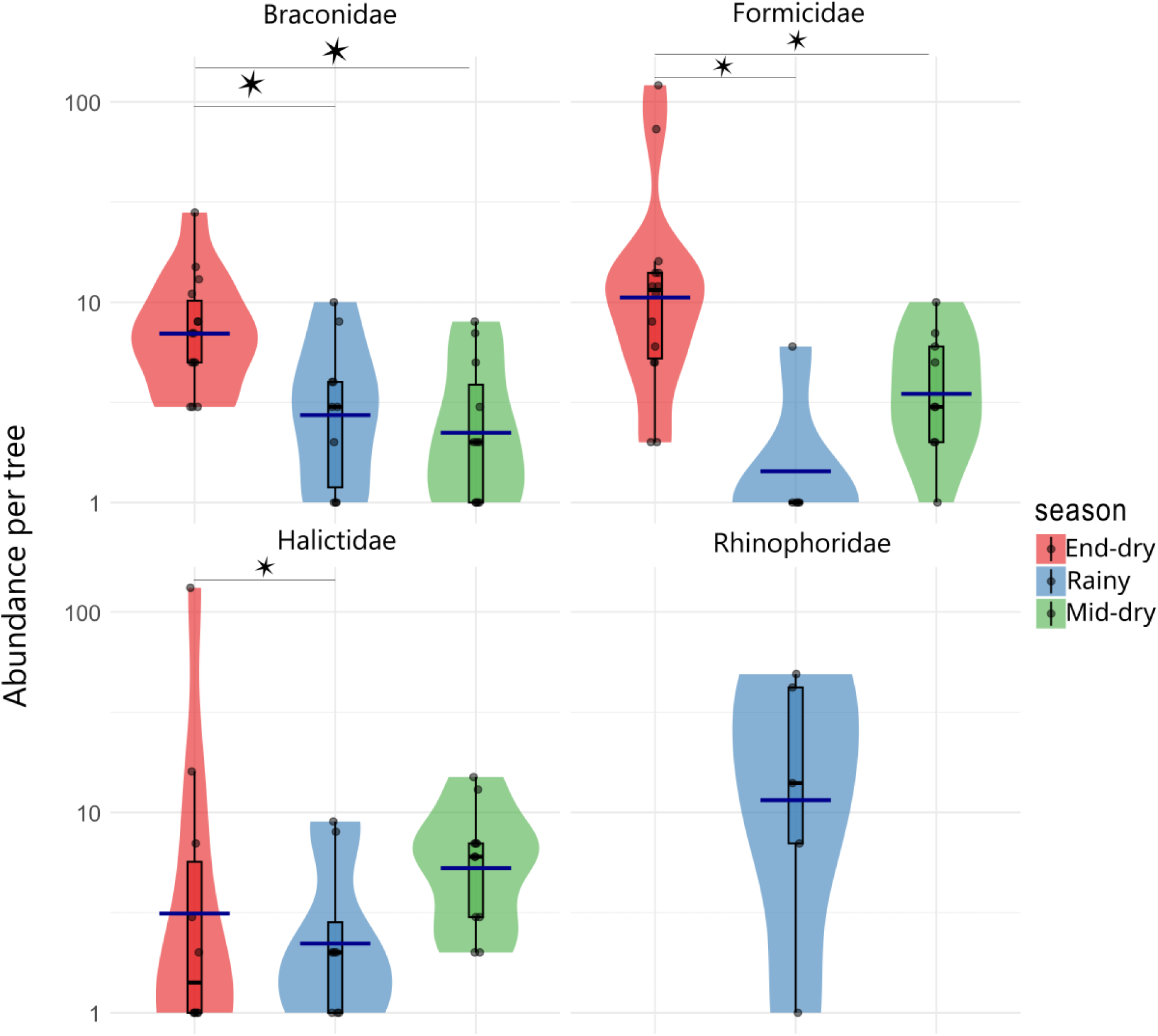
Violin plots showing the distribution across seasons of the mean abundance per tree of four families (Braconidae, Halictidae, Formicidae and Rhinophoridae) visiting flowers of *Balanites aegyptiaca* (results for the two sampling methods combined). In red: end-dry season (June); In blue: rainy season (August); in green: mid-dry season (February). Each violin represents the probability density of abundance values within a season, with individual sampling points overlaid as jittered dots to make overlapping data visible. Boxplots inside each violin show the median and interquartile range. Dark blue bars indicate the mean abundance for each group. See statistical details in Table S5. Asterisks indicate significant differences (p < 0.05). Count data were analysed using negative binomial generalized linear models with a log link. Data are presented on log₁₀-scaled axes for visualization purposes only.

##### 1. Halictidae

These bees were abundant year-round due to the consistent availability of the floral resources on which they rely. However, they were significantly more abundant at the end of the dry season (June) than in the rainy season (August; χ² = 8.44, df = 2, p = 0.015), a difference confirmed by a post-hoc pairwise comparison (p = 0.013; see Figure 9 and for statistical details Table S5).

##### 2. Rhinophoridae

These flies were completely restricted to the rainy season, Rhinophoridae were almost exclusively captured using pan traps (see Figure 9 and for statistical details Table S5).

##### 3. Braconidae

Abundance of these parasitoid wasps varied significantly among seasons (χ² = 16.31, df = 2, p < 0.001). They were significantly more abundant at the end of the dry season than in both the rainy season (p = 0.011) and the mid-dry season (p = 0.001; see Figure 9 and for statistical details Table S5). All Braconidae sampled in our study belonged to the subfamily Microgastrinae, all of which are parasitoids of diverse Lepidoptera (Whitfield et al., 2018). Their abundance coincided with a strong peak in abundance of Lepidoptera (see section 3.1. above).

##### 4. Formicidae

Ant abundance in our captures also varied significantly among seasons (χ² = 23, df = 2, p < 0.001). Ants were significantly more abundant at the end of the dry season than in both the rainy season (p < 0.001) and the mid-dry season (p < 0.001; see Figure 9 and for statistical details Table S5).

## Discussion

### (1) The surprising diversity of flower-visiting insects

The flowers of the desert date (*Balanites aegyptiaca*) provide freely accessible nectar and pollen, resources that should be exploitable by insects of various sizes, morphologies, and behaviours. This study showed that the diversity of insects visiting its flowers was in fact very high, comprising at least 371 morphospecies from 10 insect orders. For a semi-arid ecosystem near the southern edge of the Sahara, intensively grazed and browsed by livestock, this diversity was unexpectedly high. For comparison, in a study of the overall assemblage of flower visitors to plants in a more humid savanna in Kenya with less extreme seasonality, greater plant species richness, and no livestock, Guy et al. (2021) found around 300 insect morphospecies. Furthermore, estimations based on species abundance distributions suggest that the true species richness of visitors to *B. aegyptiaca* flowers in the northern Sahel is substantially higher than the observed richness. Ongoing analysis by barcoding of our morphospecies confirms our identifications.

We found that the two sampling methods we used captured somewhat different insect assemblages. We first analysed results for these two methods separately to understand these differences. Small, rapidly flying insects such as halictid bees and many flies were difficult to capture by netting but were easily captured with pan traps, whereas larger insects were more easily captured by netting and non-flying insects such as ants were easily captured by both methods. Each of these methods also has other biases (Wilson et al., 2008; Vrdoljak & Samways, 2012; Westerberg et al., 2021). Together, they capture a greater diversity of insects than either method separately (Popic et al., 2013). In further analyses, we thus combined samples from these two complementary methods to investigate global patterns.

The two most abundant and diverse orders, Hymenoptera and Diptera, accounted for a large proportion (77 %) of all individuals of flower-visiting insects. The most abundant insects at flowers were ants. Ants play diverse roles as flower visitors, consuming nectar and preying upon other floral visitors (florivores or pollinators; Romero & Vasconcellos-Neto, 2004), but for most plants they are unlikely to be important pollinators. Ants are followed in abundance by solitary bees (Halictidae). Small flies belonging to a diverse set of Diptera families were next in abundance. All of these groups showed strong seasonal variation in abundance and, for Diptera, in diversity as well.

To our knowledge, few studies have sampled the abundance and diversity of flower-visiting insects over multiple periods in such a highly seasonal environment. Despite strong logistical constraints, we sampled not only during the short rainy season but also during the middle and near the end of the dry season, and showed that each of these periods was characterised by a highly distinct insect fauna. Sampling at additional periods, for example, during the transition between the dry and rainy season when trees increase their vegetative activity (Novais et al., 2019), might yield an even finer-scale picture of seasonal variation in insect diversity.

### (2) Predominance of solitary bees, mostly Halictidae

Exclusively dependent on floral resources throughout all life stages, bees are often considered the most important of all pollinators. A striking feature of the bee fauna visiting the flowers of *B. aegyptiaca* is the virtually complete absence of eusocial bees. Only two individuals of *Apis mellifera* Linnaeus, 1758 and no meliponine bees were recorded. *Apis mellifera* is generally very rare in the region (N. Medina-Serrano, unpublished data). The bee fauna is completely dominated by solitary bees, predominantly Halictidae. Development and maintenance of colonies of eusocial bees requires long flight periods and a continuous supply of floral resources during the flight period. Seasonal scarcity of floral resources may lead to their relative rarity in dry environments like the northern Sahel (Medina-Serrano et al., 2025). Moreover, unlike tropical eusocial bees, solitary bees mainly nest in the ground (Antoine & Forrest, 2021). The abundance of ground-nesting species in the study area may be related to the rarity of soil-dwelling fungal pathogens in arid environments (Michener, 1979).

Four morphospecies of Halictidae were among the most abundant species in all surveys, with three species present in every season and at almost every site. The year-round presence of these species suggests they may have multiple generations per year. The halictid species present all belong to the subfamily Nomioidinae. Solitary and usually ground-nesting, bees of this subfamily are relatively insensitive to habitat degradation (Williams, 2011). They are the most abundant bees in the Sahara Desert and are predominant pollinators in these ecosystems (Pesenko & Pauly, 2005). Halictidae species captured in our study appear to belong to the genera *Nomioides* (morphospecies halict 1, 2, 3) and *Ceylalictus* (morphospecies halict 17). They could be effective pollinators of *B. aegyptiaca* and other trees in the ecosystem. Pesenko & Pauly (2005) describe the presence of *Ceylalictus muiri* (to which halict 17 appears to correspond) on *Acacia* trees and of *Nomioides* species on *Vachellia tortilis* subsp. *raddiana*, another dominant tree species in our study area.

### (3) Seasonal variation in bee abundance and in the availability of floral resources

Because bees rely entirely on floral resources for food in both immature and adult stages, their seasonal patterns are expected to reflect the availability of floral resources. Halictidae of subfamily Nomioidinae are polylectic, i.e., they visit flowers and consume pollen of many plant species (Pesenko & Pauly, 2005). Their abundance should thus reflect variation in abundance of overall floral resources, rather than being tied to the flowering phenology of a single tree species such as *B. aegyptiaca.* Bees were most abundant in our samples in the dry season, coinciding with the flowering period of most tree species in the region (Arbonnier, 2000). Seasonal variation in bee abundance thus indeed appears to be correlated with variation in the availability of floral resources.

Whereas flowering of herbaceous plants is restricted to the rainy season, the only period when their shallow root systems can access water, the deep roots of trees can access water year-round, permitting flowering at any time of year. Numerous tree species in the seasonally dry tropics are characterised by peak flowering in the dry season (Janzen, 1967). The most general hypothesis to explain this pattern is that vegetative competition with neighbouring trees demands maximum investment in growth during the rainy season, a small proportion of resources being stored for flowering in the dry season, when growth stops and competition is relaxed (Janzen, 1967). This hypothesis would appear not to apply in the northern Sahel, where trees are usually scattered. However, many tree species of the northern Sahel also occur in closed-canopy dry forests further south, and their phenology may have evolved in adaptation to those environments. Also, dry-season flowering has other advantages that apply not only in closed-canopy dry forest but also in thinly wooded savannas. For example, flowers of trees are an important source of water and energy for wasps, bees, flies and butterflies that pass the dry season as adults, and selective pressure to maintain pollinator populations may contribute to driving dry-season flowering peaks. Also, with greater evaporation of nectar in the dry season, bees and other insects must visit a larger number of flowers to meet their requirements for water (Janzen, 1967).

### (4) Diverse seasonal patterns in flower-visiting insects other than bees

Other flower-visiting insects depend on diverse resources, often using nectar, pollen, or both as fuel in their search for these resources. Our study revealed that the floral visitors of *B. aegyptiaca* included insects with a great diversity of biologies: predators, parasitoids, phytophagous insects, and detritivores (decomposers). The highly varied seasonal patterns in availability of the resources exploited by these insects may contribute to explaining the diverse seasonal patterns in their abundance and diversity. For example, most Hemiptera captured were Cicadellidae (leafhoppers) and other plant-sucking insects. It is widely known that insects of these groups on tree hosts are frequently dependent on young growth because it is easier for their piercing-sucking mouthparts to penetrate less-lignified tissues and access the plant’s sap, and this may explain their greater abundance in the rainy season. In studies of seasonality in dry tropical forest reviewed by Kishimoto-Yamada & Itioka (2015), leafhoppers and other plant-sucking Hemiptera were found to have peak activity in the wet season. The soft, worm-like larvae of Diptera are dependent on humid environments (Courtney et al., 2009), and this may similarly explain their greater abundance in the rainy season. Adults of Coleoptera found in our study often had thick cuticles, and this may help explain their apparently lower sensitivity to seasonal variation in precipitation and humidity. However, beetles have very diverse lifestyles, and in tropical dry forests, representatives of several families have peak activity in the rainy season (Kishimoto-Yamada & Itioka, 2015).

Ants were the insect group most frequently captured at flowers, represented by 10 morphospecies and a total of 350 individuals. They were much more abundant at flowers in June (end of the dry season) than in the two other months sampled. Three morphospecies, each belonging to a different genus, together accounted for 275 individuals (78.6%). Of these, *Monomorium* sp. (204 individuals) and *Crematogaster* sp. (21) were captured only at the end of the dry season (June). Of the 50 individuals of *Tetraponera* sp., 46 were captured in this same season. This seasonal variation probably reflects not a change in the abundance of these generalist predators and omnivores, which live in year-round and perennial colonies, but rather seasonal differences in the diversity and abundance of the different kinds of resources available to ants. A wider variety of insect prey is likely to be available in the rainy season, associated with young plant growth, whereas flowers and the insects attracted to them may be the only sources of water, energy, and protein for ants in the dry season (Rico-Gray & Oliveira, 2007).

Other flower-visiting insects are less long-lived, and seasonal patterns likely reflect real variation in their abundance and diversity. For some of these, the observed seasonal patterns can be plausibly explained by features of their biology. Diptera tended globally to be most abundant in the rainy season. As noted above, this could be related to the reliance of larval stages of Diptera of many families on moist habitats. One family of Diptera whose abundance in and limitation to the rainy season can be easily explained in terms of larval resources is the Rhinophoridae. These flies are specialist parasitoids of woodlice (Isopoda) (Wood et al., 2018). These litter-dwelling detritivorous crustaceans have gills (inherited from their marine ancestors) that must be kept moist. In xeric environments they tend to be nocturnal and are most likely to be active at the soil surface only in the rainy season (Hornung, 2011). However, one Diptera family, Mythicomyiidae, was abundant in the middle of the dry season, and captured almost exclusively in that season. We can offer no explanation for this notable exception, because very little is known about the biology of these vastly understudied small flies. Records for a few species indicate they may be parasitoids of diverse insects. Like other parasitoids, they would encounter within the host the moist conditions that dipteran larvae require. They are particularly abundant and diverse in arid environments and are frequently observed at flowers (Evenhuis & Lamas, 2017). Other various small Diptera, belonging to families Chloropidae, Phoridae, and Milichiidae, were also abundant visitors of *B. aegyptiaca* flowers. Owing to their small size, their likely small flight ranges, and their low density of body hairs, these species are unlikely to be efficient pollinators of *B. aegyptiaca*, but given their abundance their contribution to pollination may nevertheless be significant (Kirk-Spriggs & Muller, 2017).

Other insect groups, such as hymenopteran parasitoids, attack diverse life stages (eggs, larvae, pupae) of very diverse insects, each with its own annual cycle, and this is reflected by their diverse phenologies. Lack of knowledge about their biology prohibits interpretation of seasonal patterns, with one possible exception. Braconid wasps of the subfamily Microgastrinae showed a pronounced seasonal peak at the end of the dry season, about four times higher than in the two other seasons sampled. These wasps are exclusively parasitoids of diverse lepidopteran larvae (Fernandez-Triana et al., 2020). Interestingly, Lepidoptera were captured in our samples (pan-trap only) almost exclusively in this same season. Emergence of Braconidae may thus be timed to anticipate abundance of their hosts, which, as argued above, may be timed to anticipate flushes of young leaves. Emergence of adult Lepidoptera is likely timed to anticipate flushes of plant growth at the end of the dry season before the onset of rains (Ryan et al., 2017).

A fundamental problem in interpreting seasonal patterns in flower-visiting insects is that little appears to be known about how different insects in the region survive unfavourable seasons (for many, the dry season). If this knowledge exists, it has never been synthesised. Several strategies have been documented, including seasonal migration and passing the dry season as active adults (Medina-Serrano et al., 2025), but for most groups there is no available information.

### (5) Variation among sites

For several families (Diptera: Ceratopogonidae, Chloropidae, Phoridae, Rhinophoridae; Hymenoptera: Crabronidae, Formicidae, Halictidae, Platygastridae), the number of individuals captured in the Restored site (within the Koyli Alpha Community Natural Reserve) was somewhat higher than in the Unrestored site (intensively grazed and browsed rangeland near the reserve). One of these families, Rhinophoridae, was captured almost exclusively in the Restored site. However, our results tell us little about the effects of restoration actions on assemblages of flower-visiting insects, as the lack of replication prevents any conclusions. In any case, restoration actions have probably not been in place long enough to produce detectable effects. The Koyli Alpha reserve was inaugurated only seven years ago. Few studies of ecological restoration in other environments report a recovery in insect diversity before a decade or so (Suding, 2011). Working in another site within the area of GGW projects in Senegal, Brandt et al. (2020) showed that the number of trees in restored parcels was higher than in surrounding unrestored areas, but that trees in the these parcels were smaller and that tree cover was consequently lower. This suggests that tree cover will increase in the GGW parcels, with likely positive impacts on biodiversity of flower-visiting insects as trees reach reproductive maturity.

One of our sites was located in a topographical depression, where soils remain humid longer into the dry season. Such sites harbour greater density and species richness of trees and are seen as refugia for biodiversity in the face of intensive land use and climate change (Dendoncker et al., 2023). Like almost all tree species in the region, trees in these sites are insect-pollinated. If their greater tree species richness also reflects higher functional diversity, they should support greater diversity of flower-visiting insects (Kireta et al., 2024). We found surprisingly little indication that flower-visiting insects were more abundant in these tree-rich and relatively humid sites. In fact, abundance of insects per tree in the Depression site was lower than in the Restored site. Interestingly, insect abundance at a tree was negatively correlated with local tree density (as estimated by mean distance to the 10 closest neighbouring trees; Figure S5, Table S1), and local tree density was significantly higher in the Depression site than in the two other sites (Figure S5). We thus cannot separate an effect of site on insect abundance from an effect of local tree density. Curiously, insect abundance was much less variable among trees in all seasons in the Depression site than in the two other sites (Figure 6). We have no explanation for this difference. In contrast to abundance, observed species richness was as high in the Depression site as in the Restored site.

Furthermore, estimated richness (ChaoRichness) was much higher in the Depression site than in the Restored site, especially for flying insects captured by hand nets. This (along with the somewhat contrasting patterns of observed species richness and Simpson diversity between these two sites) suggests that observed richness particularly underestimates expected richness in the Depression site, underscoring the likely importance of topographical depressions in this region as a refuge not only for rare plant species (Dendoncker et al., 2023) but also for insect diversity.

The finding that insect abundance at a tree was significantly negatively correlated with local tree density is puzzling. Because we do not know if neighbouring trees were flowering in each of the sampling periods, we cannot judge whether this pattern is related to distribution of floral resources. The finding does suggest, however, that further analyses should consider larger spatial scales (e.g., Musters et al., 2021).

Comparing indicators of diversity of flowering-visiting insects across sites in the same season, we detected a marked difference only during the mid-dry season, between the Depression and the Restored sites. Across all diversity indices (Hill numbers q = 0, 1, and 2), diversity was lowest at the end of the dry season and highest during the mid-dry and rainy seasons, particularly at the Depression site. The similarity in the responses of Shannon (q = 1) and Simpson (q = 2) indices indicates that changes in diversity were mainly driven by the dominant species rather than by rare species. In other words, seasonal variation in diversity reflected overall changes in insect activity and the identity of dominant taxa, rather than shifts in the degree of dominance or evenness within assemblages.

### (6) Perspectives and open questions

Our study reveals several critical gaps in knowledge about interactions between plants and flower-visiting insects in the Sahel. These must be addressed to effectively guide the maintenance, and if necessary the restoration, of the diverse ecosystem functions performed by these components of biodiversity.

First, we need to know much more about the basic biology of flower-visiting insects in the region. What is the range of plant species with which each interacts? This study was focused on the insects visiting flowers of a single common tree species. Like *B. aegyptiaca*, flowers of other common trees of the region offer pollen and nectar freely accessible to diverse insects (Medina-Serrano et al., 2025). Studies in progress on some of these other trees show that they share many floral visitors with *B. aegyptiaca* but that assemblages also differ (N. Medina-Serrano, M. Hossaert-McKey & D. McKey, in preparation). How flower-visitor networks are structured across tree species is unknown. The herbaceous vegetation of the region, dominated by grasses (wind-pollinated), also includes numerous insect-pollinated forbs (non-graminoid herbaceous plants). Unlike trees, forbs of the region flower only during the brief rainy season. The assemblages of insects visiting their flowers are unknown. While many have ‘open-access’ flowers like those of the region’s trees, in many others morphological traits restrict access to floral rewards. These include the “bee flowers” of many faboid legumes, as well as flowers apparently adapted to hawkmoths, to settling moths, or to butterflies (Medina-Serrano et al., 2025). We do not know whether trees and forbs are associated with distinctive subnetworks of flower visitors that differ in their structure and functioning.

Halictid bees appear to be among the most important pollinators in the region. The species we found are polylectic, exploiting pollen and nectar of a broad range of flowers. The presence of each species in all seasons suggests that they are long-lived or are multivoltine, or both. Knowing their host range, life history, and behavioural ecology is key to understanding the dynamics and resilience of their populations. We do not know whether these same species also pollinate forbs in the rainy season.

The importance of Diptera as pollinators is globally underestimated (Ollerton, 2017). In Africa, Diptera reach their greatest diversity in seasonally arid environments (Kirk-Spriggs & Stuckenberg, 2010), and we recorded diverse families of small flies as abundant visitors to flowers of *B. aegyptiaca*. Little is known about the biology of many of these flies. An outstanding example is the Mythicomyiidae, tiny flies, highly diverse and vastly understudied, with numerous species awaiting description. Very little appears to be known about their reproductive ecology.

Biological investigations require accurate species identifications. The dearth of taxonomists is particularly severe in Africa, especially for difficult groups such as small Diptera. There is a great need for integrative taxonomy, combining morphology, behaviour, and molecular barcoding.

This study has produced a one-time snapshot of insect abundance and diversity at flowers of the most common tree species in the northern Sahel. Extending this snapshot to long-term monitoring will be essential for effective conservation strategies. How will climate change and continued ecosystem degradation affect insect-flower interactions? Can restoration actions carried out in initiatives such as the Great Green Wall maintain the biodiversity of flower-visiting insects? Finally, insect-flower interactions are only one facet of the interactions in which biodiversity ensures the functioning of healthy ecosystems. Each of the trees and insects we studied here is linked by other biotic interactions in interlocking networks of herbivory, predation, parasitism, seed dispersal, and the cycling of nutrients (Pilosof et al., 2017). Efforts to conserve and restore Sahelian ecosystems must embrace and understand this complexity and integrate it into management actions.

Climate change is altering insect phenology in many ways, with manifold potential consequences for interactions between insects and flowers. Whereas shifts in phenology in response to climate change are beginning to be documented in insects of temperate regions, there is a dearth of information on how insect phenology responds to climate change in the seasonal tropics. In these regions, where temperature varies relatively little across seasons, phenology is less likely to be tied to temperature and the consequences of warming for phenology are less clear (Forrest, 2016). On the other hand, changes in the amount, timing, and irregularity of rainfall are likely to have great impact on insect phenology (Janzen & Hallwachs, 2021). Careful, long-term studies of tropical insect phenology are needed to investigate correlations between seasonal activity and environmental variables that are affected by warming, such as seasonal precipitation patterns (Valtonen et al., 2013).

### (7) Limitations of the study

Our study provides the first seasonal assessment of flower-visiting insects in the northern Sahel, but several limitations should be acknowledged. First, sampling was restricted to a single locality with only three sites, which limits inference about regional-scale variation. The absence of site replication means that patterns attributed to restoration or topography should be interpreted cautiously. Second, although three contrasting seasons were sampled, temporal coverage within each was brief, and some interannual variation in flowering or insect activity may have been missed. Third, pan traps and hand-netting differ in their efficiency for different insect groups, potentially biasing estimates of relative abundance. Finally, our identification was limited to morphospecies, which restricts taxonomic resolution and prevents firm conclusions about species-level turnover. Each of the morphospecies we identified is distinct from similar species. We think it unlikely that the actual number of species in our sample is smaller than the number of morphospecies we identified. On the other hand, some morphospecies may contain cryptic species.

Future work should expand sampling across localities, integrate molecular identification, and include finer-scale environmental data to strengthen inferences about drivers of insect diversity in Sahelian ecosystems.

## Conclusion

Flowers of *B. aegyptiaca* in the intensively grazed and browsed rangelands of the northern Sahel are visited by an unexpectedly high diversity of insects. The abundance of this tree, the ease with which insects of varying size, morphology and behaviour gain access to its nectar and pollen, and its year-round flowering, combine to make it a critical resource for many insect species, which in turn play diverse roles in many other biotic interactions. The biology of insects visiting the flowers of *B. aegyptiaca*, even the most abundant ones, is very poorly known. Abundance, diversity, and species composition of assemblages of flower-visiting insects vary across seasons. As in tropical dry forests (Kishimoto-Yamada & Itioka, 2015), some insects have peak activity in the rainy season, but for others peak activity occurs in the dry season, which in the northern Sahel is exceptionally long and harsh. While some of this variation can be explained based on what we know about the functional traits of insects, our sparse knowledge of the biology and ecology of insects in the northern Sahel limits our interpretation of patterns. Knowledge of the biotic interactions that underpin ecosystem functioning, and how they vary across seasons, is crucial for monitoring, understanding, and managing the impacts of ecosystem degradation, restoration actions, and climate change on Sahelian ecosystems.

## Acknowledgments

We thank the IRL 3189 (International Research Laboratory of the CNRS, “Environnement, Santé, Sociétés”), the Institut Fondamental d’Afrique Noire Cheikh Anta Diop (IFAN Ch. A. Diop), the Département de Biologie Végétale of the Université Cheikh Anta Diop (UCAD) and their staff and students for their welcome and for organising and participating in field missions. We thank the local populations of the Ferlo, on whose land we conducted fieldwork, and the Senegalese agency responsible for land management in the region, the ASERGMV (Agence Sénégalaise de la Reforestation et de la Grande Muraille Verte). Natalia Medina-Serrano thanks Julien Haran from the Centre de Biologie pour la Gestion des Populations (CBGP) in Montpellier for all his encouragement, support and advice since the beginning of this study and for making available DNA barcoding facilities for some of the specimens collected. The authors thank the reviewers and recommender for all the suggestions that allowed us to considerably improve the manuscript.

Preprint version 4 of this article has been peer-reviewed and recommended by Peer Community In Ecology (https://doi.org/10.24072/pci.ecology.100793 ; Galloni, M. (2026) Investigating flower–insect interactions in Sahelian ecosystems: the role of *Balanites aegyptiaca*. *Peer Community in Ecology,* 100793).

## Data availability statement

The scripts of the analyses carried out and the raw data are available in Zenodo: Medina-Serrano, N. (2026). Metadata for Seasonal variation in insect assemblages at flowers of Balanites aegyptiaca, an ecologically and socially important tree species in the Ferlo region of Senegal’s Great Green Wall corridor [Data set]. Zenodo. https://doi.org/10.5281/zenodo.18959589

## Ethics approval statement

Senegal adheres to the Nagoya protocol, but no official procedure has yet been initiated. The transportation of sampled specimens was carried out with the agreement of the institutions (ASERGMV and Institut Fondamental d’Afrique Noire Cheikh Anta Diop).

## Supporting information

**Figure S1.**
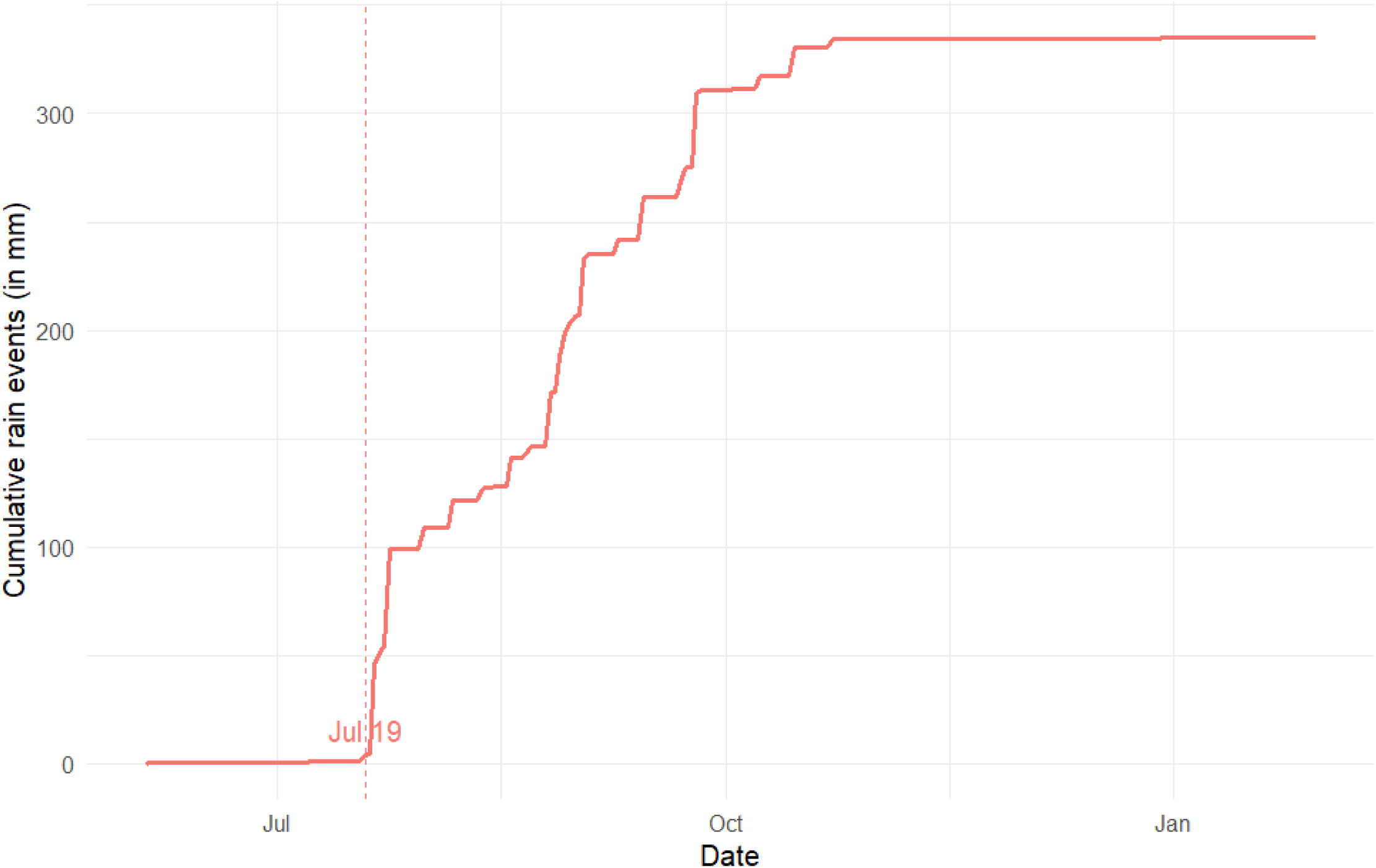
Cumulative rainfall (in mm) over time for Koyli Alpha in the year 2022, showing the first significant rain (≥ 2 mm).

**Figure S2.**
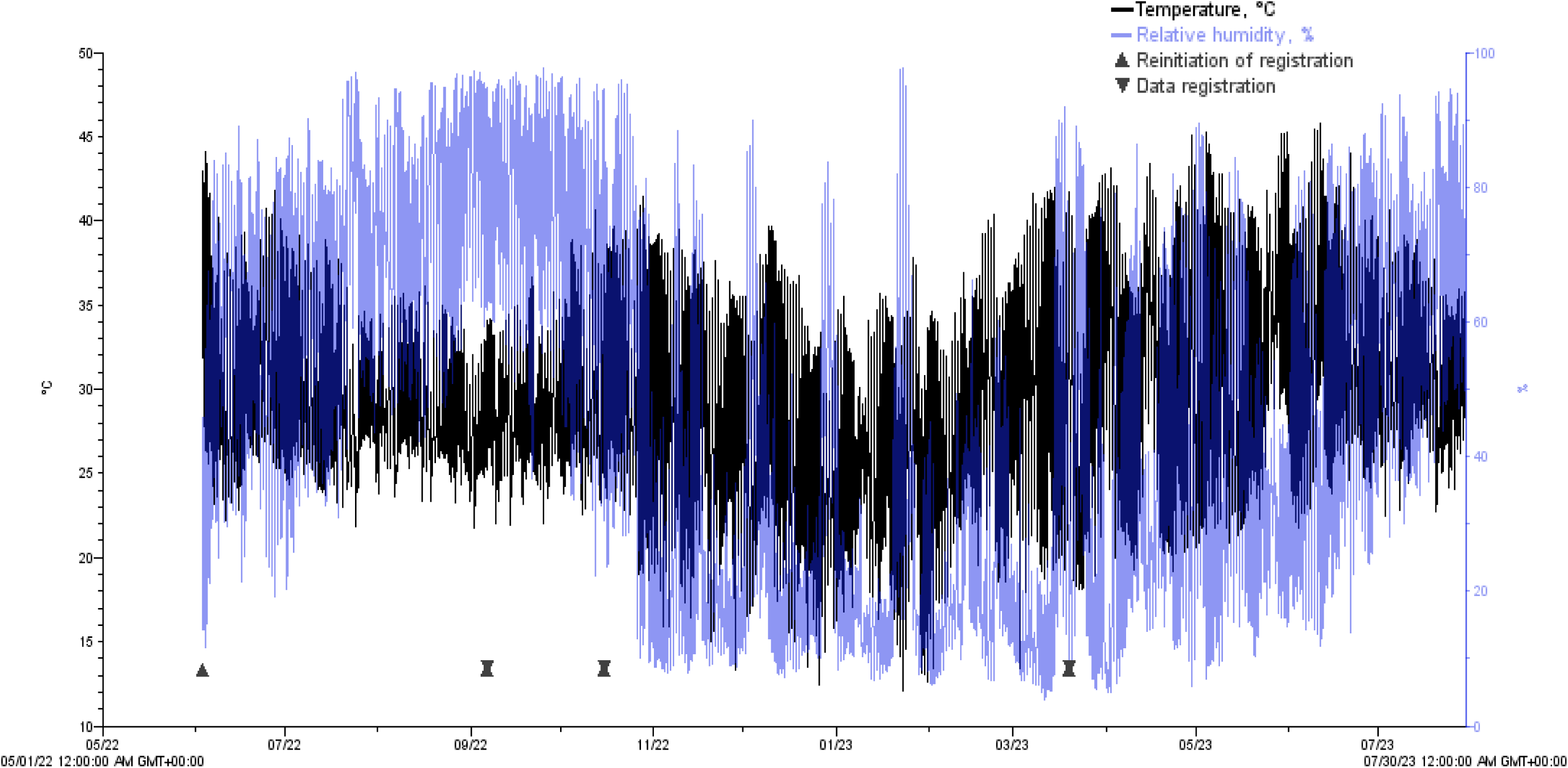
Diurnal variation in temperature and relative humidity measured at Koyli Alpha each day between 1 May 2022 and 30 July 2023. Note the strong diurnal variation in relative humidity during the dry season (November-March). Product: HOBO U23-001A Temp/RH. Onset’s HOBO Data Loggers (2018).

**Figure S3.**
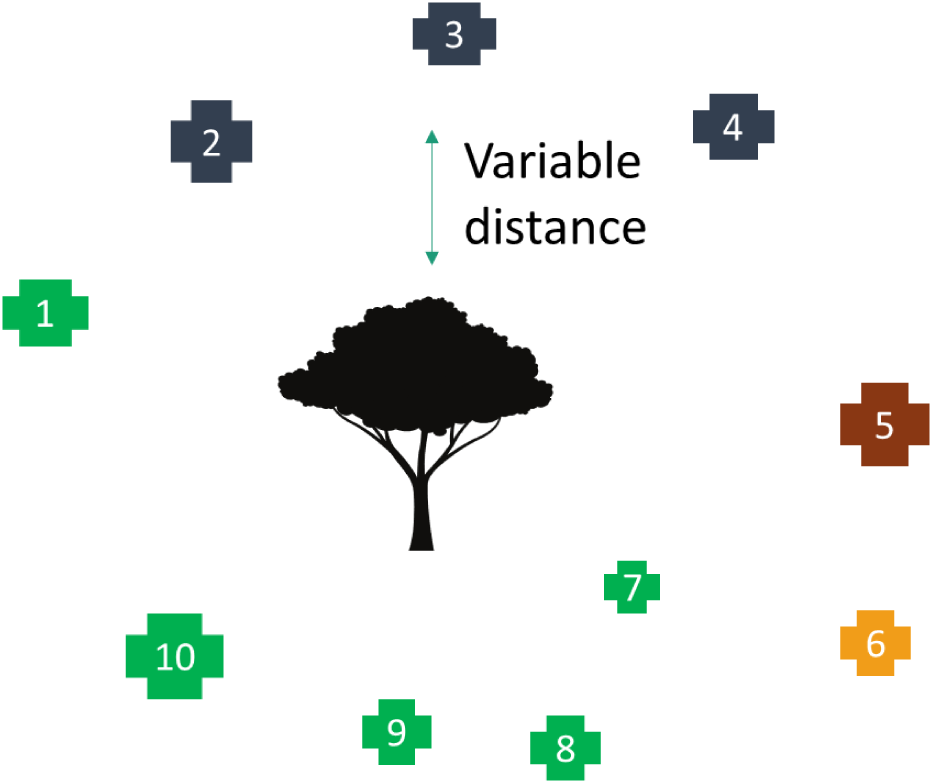
Schema showing estimation of local tree density and diversity around a sampled individual *Balanites aegyptiaca*. The 10 closest trees to each focal tree included 2-4 tree species. The crosses represent the 10 trees, and the different colours of crosses indicate different tree species. Size of the cross indicates relative size of the tree.

**Figure S4.**
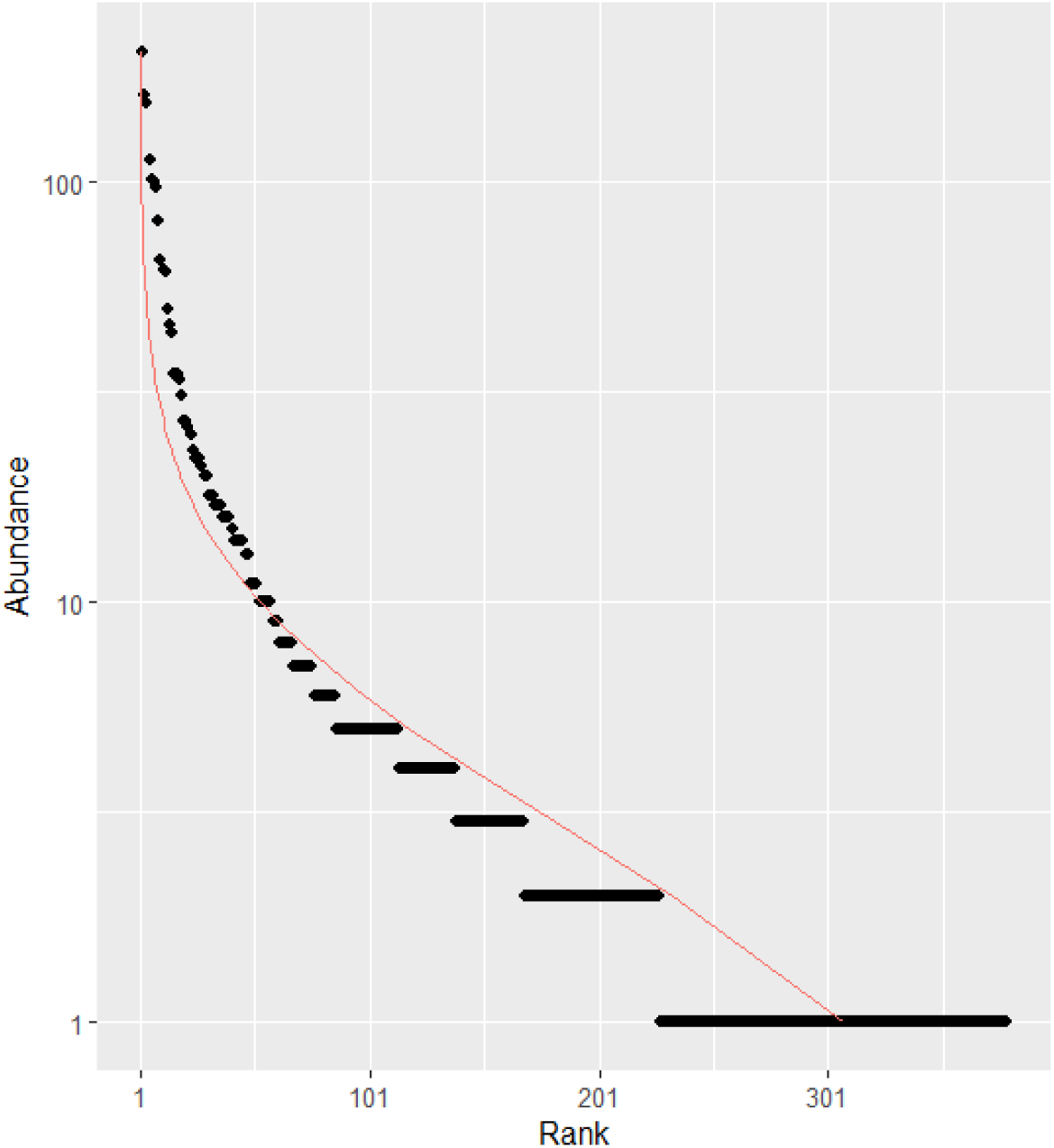
Rank-abundance curve of different morphospecies in the community of insects, combining hand net and pan-trap samples and data for all three seasons. The curve follows a log-normal distribution. The curve was done using the package entropart (Marcon & Hérault, 2015).

**Figure S5.**
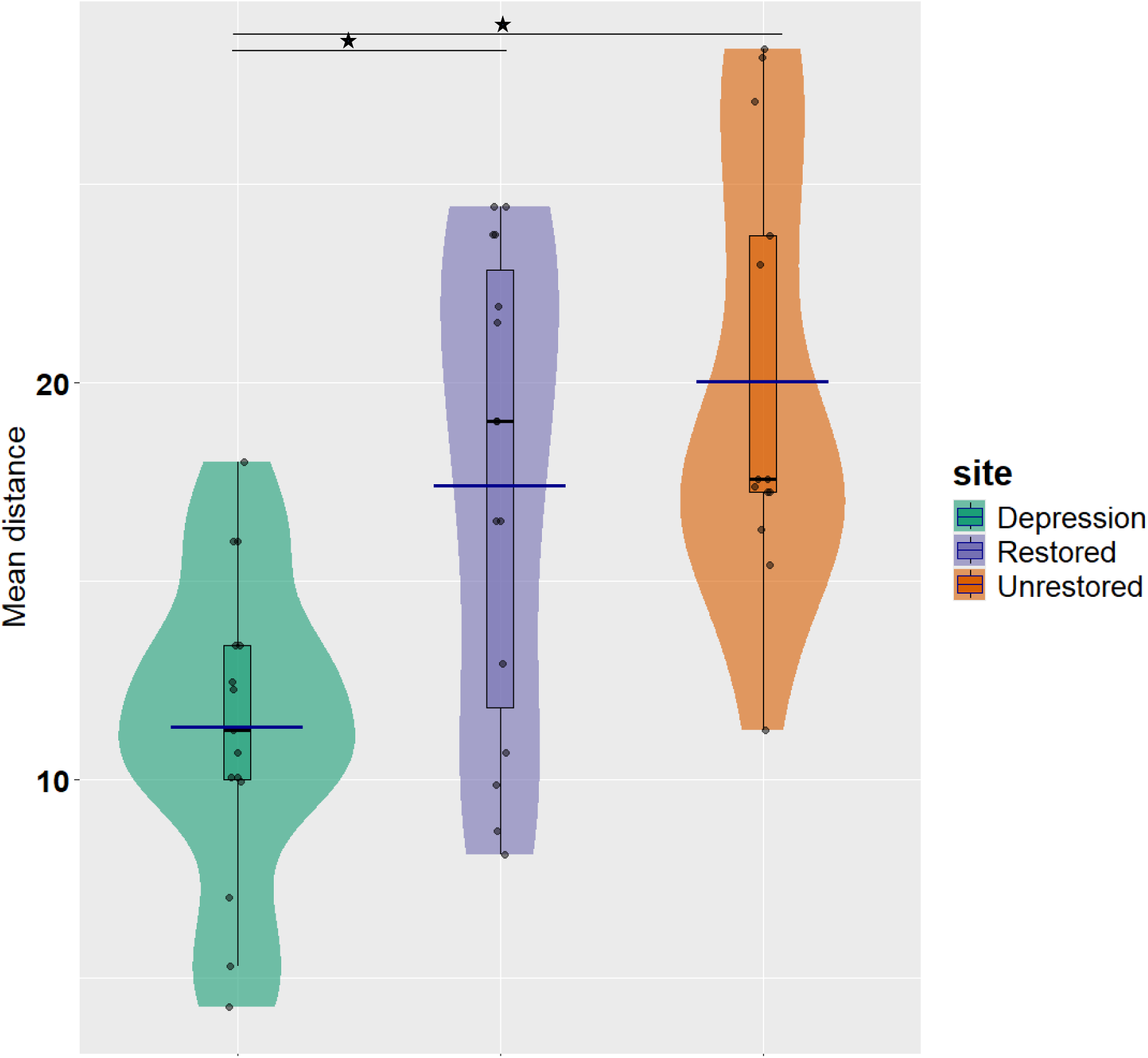
Violin plots showing the mean distance to the 10 closest trees for each site. Each violin illustrates the probability density of values within a site, with individual sampling points overlaid as jittered dots to reduce overlap. Boxplots inside each violin show the median and interquartile range. Dark blue horizontal bars represent group means. Colours indicate site categories (green = Depression, purple = Restored, orange = Unrestored).

**Figure S6.**
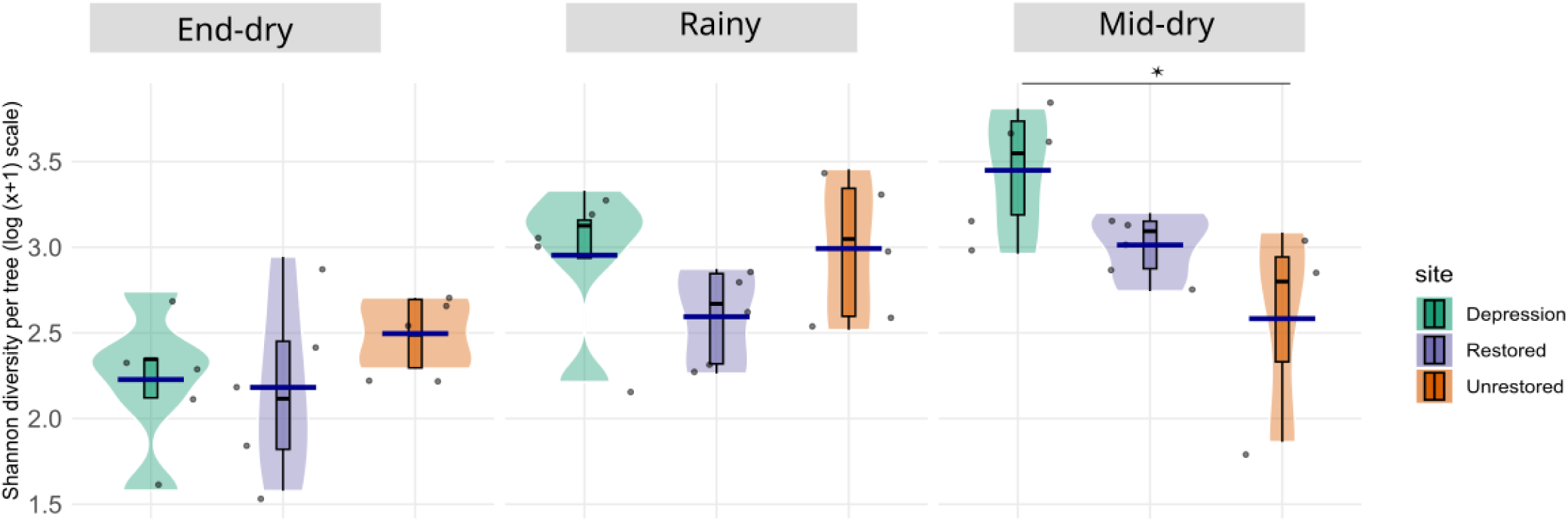
Violin plots of insect assemblages at flowers of *Balanites aegyptiaca*, showing for each site log1p(Shannon diversity index (q=1) per tree) across seasons: end-dry (June), rainy (August), and mid-dry (February). Each violin illustrates the probability density of values within a season, with individual sampling points overlaid as jittered dots to reduce overlap. Boxplots inside each violin show the median and interquartile range. Dark blue horizontal bars represent group means. Colours indicate site categories (green = Depression, purple = Restored, orange = Unrestored), and facets allow comparisons across seasons. Shannon diversity (q=1) was significantly affected by season (R²_adj = 0.454, F = 5.373, df = 8,34, p = 0.00021). ANOVA showed a significant main effect of season (p = 0.00012), while site (p = 0.411) and site × season interaction (p = 0.055) were not significant. Pairwise Tukey comparisons revealed that during the mid-dry season, Shannon diversity was significantly higher in the Depression site compared to the Unrestored site (p = 0.014). These findings highlight seasonal changes in diversity, with notable differences among sites emerging in the mid-dry season.

**Table S1.** Results of the two different Generalised Linear models used to explore the effect of season, mean distance to the 10 closest trees and the % canopy flowering of the sampled tree on insect abundance (insects from the two sampling methods were grouped).

| Model | Formula | AIC | Lognormal R<sup>2</sup> | Predictor | Likelihood Ratio $\chi^2$ | df | p-value |
| --- | --- | --- | --- | --- | --- | --- | --- |
| **Generalised Linear Model** | Abundance ~ season+ flowering + mean distance | 442.28 | 0.2119 | Season | 1.26 | 2 | 0.5330 |
|  |  |  |  | Flowering | 1.16 | 1 | 0.2819 |
|  |  |  |  | Mean distance | 5.81 | 1 | **0.0160** |
| **Generalised Linear Model** | Abundance ~ site + flowering + mean distance | 439.01 | 0.2589 | Site | 4.7332 | 2 | 0.0938 |
|  |  |  |  | Flowering | 1.4053 | 1 | 0.2358 |
|  |  |  |  | Mean distance | 1.5089 | 1 | 0.2193 |
| **Generalised Linear Model** | Abundance ~ site*season | 441.1 | 0.3730 | Site x Season | 9.25 | 4 | 0.0551 |
|  |  |  |  | Season | 5.53 | 2 | 0.0630 |
|  |  |  |  | Site | 14.01 | 2 | **0.0009** |

**Table S2.** Tukey-adjusted pairwise comparisons for insect abundance (negative binomial GLM) for the model *Abundance ∼ site*season*. (All estimates on the log scale; p-values Tukey-adjusted.) Pairwise seasonal comparisons within (Tukey-adjusted) revealed that the only significant seasonal effect occurred at the Depression site, where end-dry season differed significantly from mid-dry season (estimate = –0.8900, *z* = –2.38, p = 0.0456). This contrast indicated lower values in end-dry season relative to mid-dry season. No other seasonal contrasts at this site were significant. Similarly, no significant seasonal differences were detected at either the Restored or Unrestored sites, as all corresponding p-values >0.23. Therefore the detailed results are not included in this table.

| Subset | Conditioning level | Contrast | Estimate | SE | z | p-value |
| --- | --- | --- | --- | --- | --- | --- |
| Sites (averaged over seasons) | All seasons | Depression – Restored | -0.7020 | 0.213 | -3.291 | **0.0029** |
|  |  | Depression – Unrestored | -0.4940 | 0.225 | -2.196 | 0.0719 |
|  |  | Restored – Unrestored | 0.2080 | 0.223 | 0.932 | 0.6200 |
| Sites within season | End dry (June) | Depression – Restored | -1.4293 | 0.372 | -3.840 | **0.0004** |
|  |  | Depression – Unrestored | -1.0832 | 0.373 | -2.902 | **0.0104** |
|  |  | Restored – Unrestored | 0.3460 | 0.365 | 0.947 | 0.6104 |
|  | Rainy (August) | Depression – Restored | -0.7315 | 0.368 | -1.987 | 0.1153 |
|  |  | Depression – Unrestored | -0.1603 | 0.370 | -0.433 | 0.9018 |
|  |  | Restored – Unrestored | 0.5712 | 0.367 | 1.555 | 0.2652 |
|  | Mid-dry (February) | Depression – Restored | 0.0548 | 0.368 | 0.149 | 0.9879 |
|  |  | Depression – Unrestored | -0.2387 | 0.423 | -0.564 | 0.8394 |
|  |  | Restored – Unrestored | -0.2935 | 0.424 | -0.693 | 0.7678 |

**Table S3.**
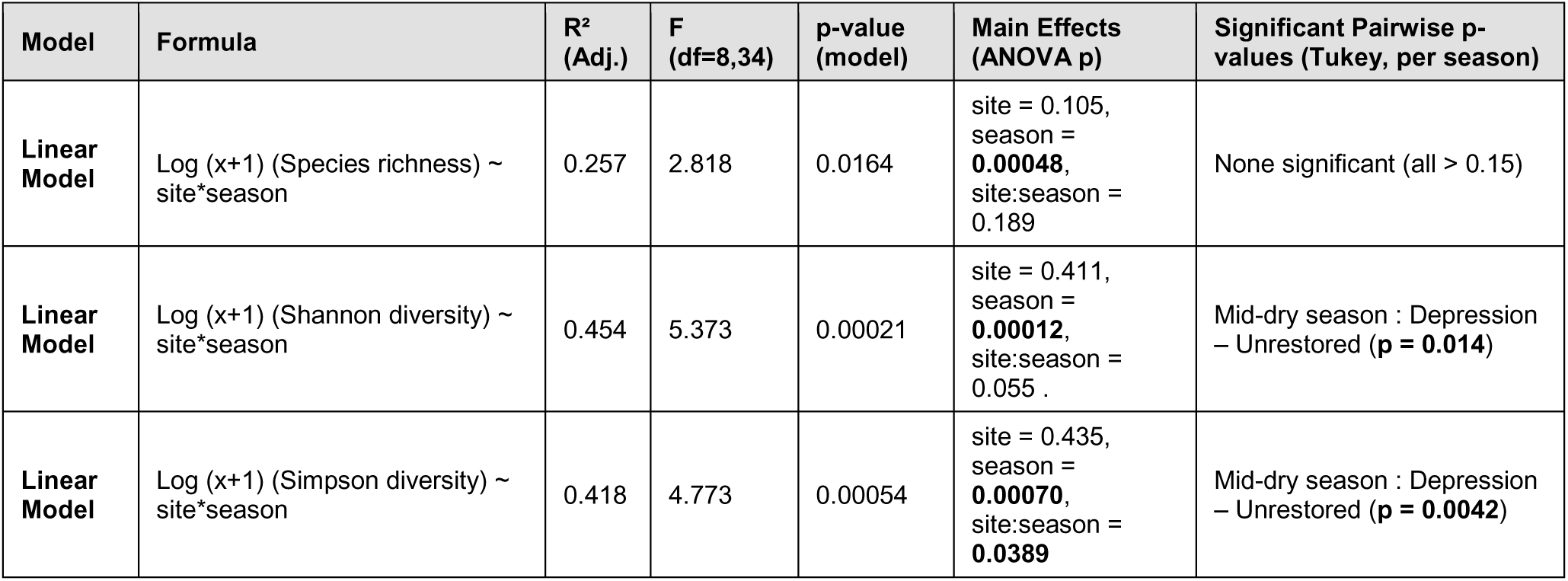
Results of the models (Generalised Linear Model, Linear Model and Linear Mixed Model) used to explore the effect of season (End-dry, Rainy, Mid-dry) in interaction with the effect of site (Depression, Restored, Unrestored) on species richness, Shannon diversity, Simpson diversity index (combining results from the two sampling methods).

**Table S4.** Results of the Generalised Linear models comparing the seasonal effects for Hymenoptera, Diptera, Coleoptera and Hemiptera (same model, testing the different order) and insects from the two sampling methods were grouped.

| Model | Formula | Lognormal R<sup>2</sup> | Analysis of variance | Pairwise comparisons (p-values) |
| --- | --- | --- | --- | --- |
| **Generalised Linear Model** | Hymenoptera abundance ~ season | 0.3247 | LR $\chi^2 = 15.02$, df = 2, p = **0.0005** | Rainy - Mid-dry: **0.0360**,<br>Rainy - End-dry: **0.0002**,<br>Mid-dry - End-dry: 0.3748 |
| **Generalised Linear Model** | Diptera abundance ~ season | 0.3391 | LR $\chi^2 = 12.95$, df = 2, p = **0.0015** | Rainy - Mid-dry: 0.6839,<br>Rainy - End-dry: **0.0007**,<br>Mid-dry - End-dry: **0.0166** |
| **Generalised Linear Model** | Coleoptera abundance ~ season | 0.1140 | LR $\chi^2 = 3.80$, df = 2, p = 0.1497 | Rainy - Mid-dry: 0.2044,<br>Rainy - End-dry: 0.9937,<br>Mid-dry - End-dry: 0.1706 |
| **Generalised Linear Model** | Hemiptera abundance ~ season | 0.4614 | LR $\chi^2 = 15.82$, df = 2, p = **0.0004** | Rainy - Mid-dry: 0.7709,<br>Rainy - End-dry: **0.0002**,<br>Mid-dry - End-dry: **0.0095** |

**Table S5.** Results of the Generalised Linear models comparing effects of season for Braconidae, Halictidae and Formicidae, grouping insects from both sampling methods. No statistical test was possible for Rhinophoridae, which were captured exclusively during the rainy season (see text).

| Model | Formula | Lognormal R<sup>2</sup> | Analysis of variance | Pairwise Comparisons (p-values) |
| --- | --- | --- | --- | --- |
| **Generalised Linear Model** | Formicidae abundance ~ season | 0.6024 | LR $\chi^2 = 23$, df = 2, p < **0.0001** | Rainy - Mid-dry: 0.427, Rainy - End-dry: **0.0001**, Mid-dry - End-dry: **0.0005** |
| **Generalised Linear Model** | Halictidae abundance ~ season | 0.3332 | LR $\chi^2 = 8.44$, df = 2, p = **0.0147** | Rainy - Mid-dry: 0.450, Rainy - End-dry: 0.0125, Mid-dry - End-dry: 0.156 |
| **Generalised Linear Model** | Braconidae abundance ~ season | 0.3713 | LR $\chi^2 = 16.31$, df = 2, p = **0.0003** | Rainy - Mid-dry: 0.810, Rainy - End-dry: **0.0113**, Mid-dry - End-dry: **0.0010** |

